# Asynchronous glutamate exocytosis is enhanced in low release probability synapses and is widely dispersed across the active zone

**DOI:** 10.1101/2021.05.04.441792

**Authors:** Philipe R. F. Mendonça, Erica Tagliatti, Helen Langley, Dimitrios Kotzadimitriou, Criseida G. Zamora-Chimal, Yulia Timofeeva, Kirill E. Volynski

## Abstract

The balance between fast synchronous and delayed asynchronous release of neurotransmitters has a major role in defining computational properties of neuronal synapses and regulation of neuronal network activity. However, how it is tuned at the single synapse level remains poorly understood. Here, using the fluorescent glutamate sensor SF-iGluSnFR, we image quantal vesicular release in tens to hundreds of individual synaptic outputs (presynaptic boutons) from single pyramidal cells in culture with 4 millisecond temporal resolution, and localise vesicular release sites with ~ 75 nm spatial resolution. We find that the ratio between synchronous and asynchronous synaptic vesicle exocytosis varies extensively among presynaptic boutons supplied by the same axon, and that asynchronous release fraction is elevated in parallel with short-term facilitation at synapses with low release probability. We further demonstrate that asynchronous exocytosis sites are more widely distributed within the presynaptic release area than synchronous sites. These findings are consistent with a model in which functional presynaptic properties are regulated via a synapse-specific adjustment of the coupling distance between presynaptic Ca^2+^ channels and release-ready synaptic vesicles. Together our results reveal a universal relationship between the two major functional properties of synapses – the timing and the overall probability of neurotransmitter release.

## Main text

Synaptic transmission provides the basis for neuronal communication. When an action-potential propagates through the axonal arbour, it activates voltage-gated Ca^2+^ channels (VGCCs) located in the vicinity of release-ready synaptic vesicles docked at the presynaptic active zone^1^. Ca^2+^ ions enter the presynaptic terminal and activate the vesicular Ca^2+^ sensor Synaptotagmin 1 (Syt1, or its isoforms Syt2 and Syt9), thus triggering exocytosis of synaptic vesicles filled with neurotransmitter molecules. Neurotransmitter diffuses across the synaptic cleft, binds postsynaptic receptors and evokes further electrical or chemical signalling in the postsynaptic target cell. This whole process occurs on a time scale of a few milliseconds. Recent data demonstrate that such speed and precision are in large part achieved via the formation of nanocomplexes that include presynaptic VGCCs, vesicles belonging to a readily releasable pool (RRP) and postsynaptic neurotransmitter receptors^2–4^.

In addition to fast, synchronous release, which keeps pace with action potentials, many synapses also exhibit delayed asynchronous release that persists for tens to hundreds of milliseconds^1,5^. Asynchronous release is potentiated during repetitive presynaptic firing and is triggered via activation of multiple sensors with both low (*e.g.* Syt1) and high (*e.g.* Syt7) Ca^2+^ affinity^6^. Accumulating evidence demonstrates that the balance between synchronous and asynchronous release plays an important role in coordinating activity within neuronal networks, for example, by increasing the probability of postsynaptic cell firing and/or modulating action potential precision^7–10^. It is well established that asynchronous release levels vary among different types of neurons^1, 10, 11^. Interestingly, recent data show that the ratio between asynchronous and synchronous release can also be differentially regulated among presynaptic boutons supplied by the same axon and depends on the identity of the postsynaptic cell, which contributes to target cell-specific communication in the brain^7, 8^.

The mechanisms that control the relative contributions of synchronous and asynchronous release at the level of single synapses are however poorly understood. Variability in asynchronous release among different neuronal types has been attributed to differences in synaptic morphology (*e.g.* the coupling distance between RRP vesicles and VGCCs)^10^ and to cell-type specific expression of components of the vesicular release machinery (*e.g.* the Ca^2+^ sensors Syt1 and Syt7)^11^. Whether the same mechanisms contribute to the regulation of release modes among presynaptic boutons located on the same axon remains unclear. It is also unknown whether synchronous and asynchronous release occur from the same or different pools of synaptic vesicles. Recent functional electron microscopy analysis combined with fast high-pressure freezing have helped to visualise loci of either synchronous or asynchronous release events within a single active zone^4, 12^. Application of this approach to synaptic populations indicates that synchronous and asynchronous release events tend to occur in different sub-domains of the active zone. However, how synchronous and asynchronous release sites are located with respect to each other within the same active zone has not yet been established.

To address these questions, we developed a novel imaging technique and data analysis framework, which allowed us to directly investigate the relationship between synchronous and asynchronous glutamate release both at the level of individual active zones and across large populations of synaptic outputs from a single pyramidal neuron. Our approach (Fig. 1) is based on expression of the fluorescent glutamate reporter SF-iGluSnFR on the axonal membrane, which allows detection of glutamate release from individual synaptic vesicles with millisecond resolution^13–16^. We sparsely transfected neocortical neurons in culture with SF-iGluSnFR and established a whole-cell voltage-clamp recording in a neuron with pyramidal morphology expressing the sensor (Fig. 1A). We next imaged action potential-evoked glutamate release by monitoring changes in SF-iGluSnFR fluorescence (at a rate of 4 ms/frame) in tens to hundreds of individual presynaptic boutons supplied by the axon of the selected neuron, in response to a 5 Hz train of 51 action potentials triggered by brief somatic voltage steps (Fig. 1B-E, Methods). By applying a set of spatial-temporal filters and calculating the maximal projection of the resulting image stack, we could identify all active boutons irrespective of their release probability, as long as they released at least one vesicle during the train (Fig. 1D and Movie S1). Evoked synaptic SF-iGluSnFR responses had a stereotypical waveform with a quasi-instantaneous rising phase that was followed by a slower exponential decay, corresponding to glutamate unbinding from SF-iGluSnFR molecules (*τ* ~ 68 ms, Fig. 1E and Fig. S1)^14^. We applied a deconvolution procedure (using the averaged SF-iGluSnFR response waveform) to determine the precise timings and the amplitudes of release events (Fig. 1E, Figs. S1–S3 and Methods). The histograms of deconvolved event amplitudes at individual boutons typically showed between one and four discernible quantal peaks, consistent with the exocytosis of one or more vesicles of glutamate (Fig. 1E and Figs. S4 and S5)^13, 15, 17^. By fitting the histogram with a sum of Gaussian functions we estimated the amplitude of the SF-iGluSnFR signal corresponding to the release of a single quantum (*q*, Fig. 1E and Methods). Critically, application of this quantal analysis allowed us to estimate the number of vesicles released during each event and therefore directly compare vesicular exocytosis among different synapses. We note that the absolute values of the normalised evoked SF-iGluSnFR signals (Δ*F*/*F*_0_) depend not only on the number of exocytosed vesicles but also on the presynaptic bouton size and geometry, which significantly vary among synapses (Fig. 1E, Figs.S4 and S5). Therefore, the use of Δ*F*/*F*_0_ is not optimal for quantitative comparison of vesicular release parameters.

**Fig. 1.**
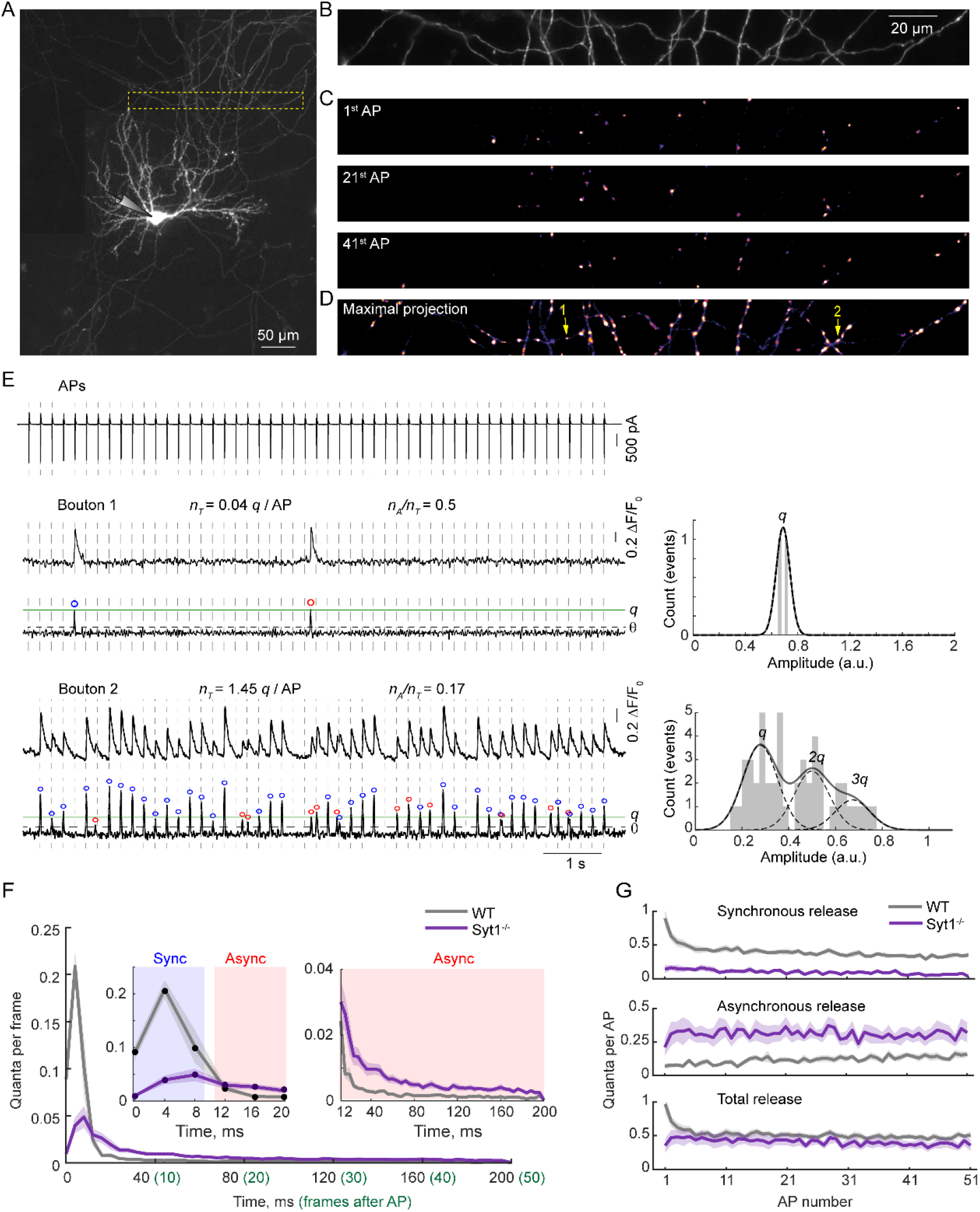
SF-iGluSnFR fluorescence imaging of quantal synchronous and asynchronous release in tens of individual presynaptic boutons supplied by a single axon. **(A** to **D)** A detailed illustration of a typical experiment. (A) Reconstructed mosaic image of a pyramidal neuron expressing the SF-iGluSnFR probe. (B) A region of interest (ROI) in the axonal arbour selected for imaging of presynaptic SF-iGluSnFR responses corresponding to the yellow box in (A). (C) Heat maps of SF-iGluSnFR responses to the 1^st^, 21^st^, and 41^st^ action potentials (APs) during 5 Hz train of 51 APs. The images are averages of 3 frames from a band-pass filtered image stack immediately after each action potential (see Methods and Movie S1). (D) Maximal projection of the band-pass filtered image stack revealing locations of all presynaptic boutons that released at least one vesicle during the stimulation train. (**E**) Analysis of quantal SF-iGluSnFR responses in two representative boutons (Boutons 1 and 2 in D). Left, somatic action potential escape currents time-aligned with band-pass filtered and deconvolved SF-iGluSnFR signals. Quantal release events were identified as local maxima on the deconvolved traces located above the threshold *θ* (horizontal dashed lines) corresponding to 4 standard deviations of the background noise (see Figs. S2, S3 and Methods). Right, quantal analysis. To determine the amplitude of SF-iGluSnFR signal corresponding to release of a single vesicle (*q*, green lines on deconvolved traces) the positions of peaks on the amplitude histograms were fitted with a sum of 4 Gaussian functions (black lines). This was then used to calculate in each bouton the total number of vesicles released per action potential (release rate *n_T_*) and the fraction of asynchronous release events (*n_A_*/*n_T_*) (see below and also Figs. S4, S5 for more examples). (**F**) Comparison of evoked vesicular release kinetics in wild type (WT) and Syt1^-/-^ neurons. The traces were obtained by first averaging in individual boutons the number of quanta detected in individual frames following each of 51 action potentials in the train (frames 0 to 50, 4 ms/frame). This was followed by calculating the cell-averaged responses (range 14 – 162 boutons per cell) and finally the mean responses across all recoded neurons. Based on the distinct biphasic time-course of vesicular release in wild type neurons, the time threshold between synchronous and asynchronous release events was set at ~10 ms after the somatic action potential (*i.e*. at the border between 2^nd^ and 3^rd^ frames, see main text for details). Using this criterion, we then classified individual release events either as synchronous (blue circles on the deconvolved traces) or asynchronous (red circles). (**G**) Comparison of synchronous, asynchronous and total release in wild type and Syt1^-/-^ neurons during the 5 Hz stimulation train. Traces represent the average number of vesicular quanta released at each action potential in a single presynaptic bouton (mean ± SEM). Wild type, N = 16 cells; Syt1^-/-^, N = 11 cells.

To define a time threshold for separation of synchronous and asynchronous release events, we calculated the average time course of vesicular exocytosis following an action potential across all recorded cells. This allowed us to compare the imaging data at single synapses to previous electrophysiological recordings from synaptic populations (*e.g.* refs.^5, 10, 18^). In agreement with the electrophysiological data, the vesicular release time-course determined with SF-iGluSnFR followed a well-defined biphasic shape (Fig.1E). We note that although synchronous release is expected to occur within 1-2 ms after the presynaptic spike^1^, due to the finite speed of action potential propagation, the onset of synchronous release is delayed by ~2-6 ms with respect to the somatic action potential (as boutons are located several hundred micrometres away from the soma, Fig. 1A)^19^. This blurs the distribution of detected synchronous release events and therefore we have used 10 ms as a threshold for the separation of synchronous and asynchronous exocytosis components.

We next tested how synchronous and asynchronous release events are distributed among synaptic outputs of individual pyramidal neurons. By using the defined 10 ms threshold we classified events as synchronous or asynchronous (blue and red colour codes respectively in Fig. 1E and subsequent figures) and calculated the total release rate (*n_T_*, average number of vesicles released per action potential) and the fraction of asynchronous release events (*n_A_*/*n_T_*) for each bouton. In line with previous studies (*e.g*. refs.^15, 20^), *n_T_* (which is directly related to the overall release probability *P_rel_*) varied widely among presynaptic boutons supplied by the same axon. Unexpectedly, we found an inverse relationship between *n_T_* and the asynchronous release fraction, which was elevated at boutons with low release probability (Fig. 2A and examples in Fig. S4). To quantify this phenomenon, we divided boutons in each recorded cell into two groups with low and high *P_rel_* (using the median value for *n_T_*). We found that asynchronous release fraction was on average ~ 1.5-fold higher in low *P_rel_* than in high *P_rel_* boutons (Fig. 2A, *n_A_*/*n_T_* = 0.31±0.04 and *n_A_*/*n_T_* = 0.20±0.02 respectively).

**Fig. 2.**
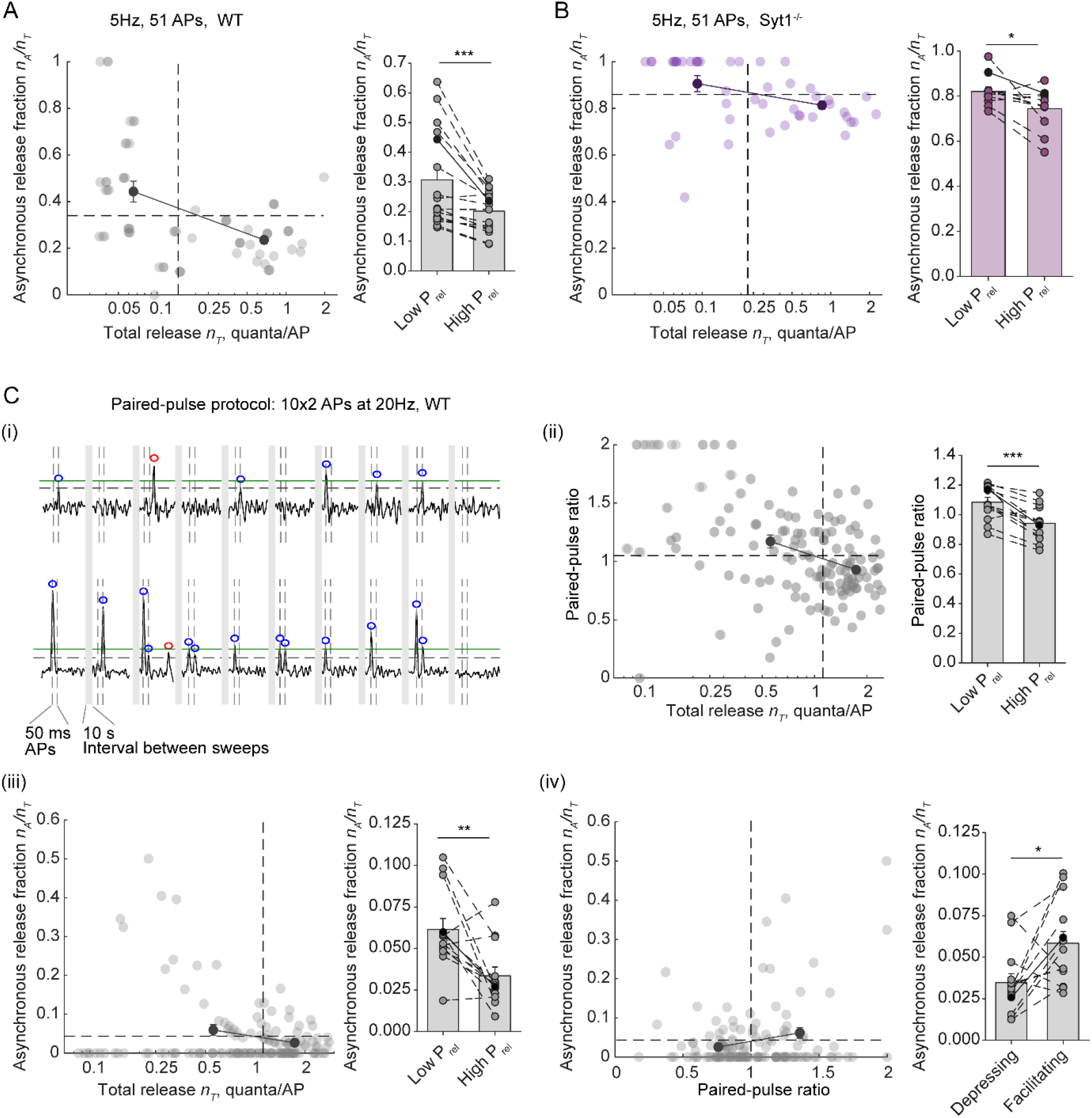
Asynchronous release is elevated in parallel with short-term facilitation in low release probability synapses. (**A**, **B**) Analysis of synchronous and asynchronous release heterogeneity among presynaptic outputs of individual wild type (A) and Syt1^-/-^ (B) neurons during 5 Hz train of 51 action potentials. Left panels, the relationship between asynchronous release fraction *n_A_*/*n_T_* and the overall vesicular release rate *n_T_* among boutons in a representative wild type (WT, n = 68 boutons) and a Syt1^-/-^ (n = 46 boutons) neurons. Horizontal dashed lines depict the average fractions of asynchronous release in each cell. Vertical dashed lines mark the median values for *n_T_*, which were used as thresholds to split boutons into groups with low and high *P_rel_*. Black dots with error bars show the average values of *n_A_*/*n_T_* in low and high *P_rel_* groups. Right, bar and dot summary plots showing increase of asynchronous release fraction in low *P_rel_* synapses across recorded cells (N = 16 wild type and N = 11 Syt1^-/-^ neurons). Data from individual cells are connected by dashed lines, black points – data from the example cells. (**C**) Analysis of the relationship between short-term plasticity and synchronicity of glutamate release rate among synaptic outputs of single neurons. (i) Paired-pulse stimulation paradigm and synaptic responses recoded in 2 representative boutons. As in Fig.1 horizontal green lines show the amplitude of SF-iGluSnFR signal corresponding to release of a single vesicle quanta (1q). Vertical dashed lines depict action potential timings. Grey boxes, 10 s inter-sweep interval. Blue and red circles mark synchronous and asynchronous release events respectively. (ii – iv) The relationships between (ii) paired pulse ratio (*PPR*) and the overall release rated *n_T_* (*PPR* in each bouton was estimated as *N_s_*_2_/(*N*_*s*1_ + *N*_*s*2_), where *N*_*S*1_ and *N*_*S*2_ are total numbers of synchronously released quanta at first and second action potentials), (iii) asynchronous release fraction *n_A_*/*n_T_* and *n_T_*, and (iv) *n_A_*/*n_T_* and *PPR* in a representative cell (n = 138 boutons, left panels), and across recorded cells (bar and dots panels, N= 13 cells). Horizontal dashed lines depict the average values of *PPR* (ii) and *n_A_*/*n_T_* (iii and iv). Vertical dashed lines in (ii) and (iii) depict the median value of *n_T_* in the example cell. Vertical line in (iv) depicts the threshold (*PPR* =1) between facilitating and depressing synapses. *P < 0.05, **P < 0.01 and *** P < 0.001 Paired t-test. To allow visual representation of the overlapping data points, the x coordinate of each point in the example cell plots shown in this figure was randomly shifted within ± 0.01 range.

It has been demonstrated that the variability in expression levels of the Ca^2+^ sensor for synchronous vesicular release Syt1 has a major role in the regulation of the balance between synchronous and asynchronous release among different types of neurons^11^. To test for a possible role of Syt1 in controlling the heterogeneity of vesicular release modes among synaptic outputs of a single neuron we repeated our experiments in neuronal cultures from Syt1^-/-^ mice. As expected, deletion of Syt1 resulted in ~ 4-5-fold decrease of synchronous and correspondingly ~ 4-5-fold increase of asynchronous vesicular exocytosis (Fig. 1F, G and Figs S4 and S5)^18^. However, in spite of the overall increase of asynchronous release, we still observed the reciprocal relationship between asyncrnonous release fraction and *P_rel_* (Fig. 2B, *n_A_*/*n_T_* =0.82±0.02 in low *P_rel_* and *n_A_*/*n_T_* = 0.74±0.03 in high *P_rel_* boutons). This indicates that the balance between different release modes at synapses supplied by a single neuron is at least in part determined by mechanisms distinct from the possible variation in the expression levels of Syt1.

Typically, synapses with low *P_rel_* display short-term facilitation of synchronous release during repetitive stimulation. Our finding that asynchronous release fraction is higher at low *P_rel_* synapses suggests that the balance between different release modes could be universally linked to the type of short-term plasticity expressed in the same presynaptic terminals. To test this hypothesis, we imaged vesicular release in response to 10 pairs of action potentials delivered at 20 Hz (50 ms inter-spike interval, 10 sec between paired pulses) and calculated in each recorded bouton the paired-pulse ratio (*PPR*), the asynchronous release fraction *n_A_*/*n_T_* and the total release rate *n_T_*. We then compared the distributions of these major functional presynaptic properties among synaptic outputs of individual neurons and found that asynchronous release fraction was indeed ~ 2-fold higher in facilitating than in depressing synapses (Fig. 2C, *n_A_*/*n_T_* = 0.058±0.007 and *n_A_*/*n_T_* = 0.034±0.005 respectively; note that as asynchronous release is potentiated during trains of action potentials^1^ the overall fraction of asynchronous release measured after two pulses was several fold lower than the fraction of asynchronous release during 5 Hz stimulation).

We next asked what are the spatial distributions of synchronous and asynchronous events within individual active zones. During vesicular exocytosis glutamate can be assumed to be released from a point source. Therefore, fusion of a single vesicle is expected to generate a bell-shaped SF-iGluSnFR fluorescence profile centred at the exocytosis site, which can be fitted using a 2D Gaussian function to determine the location of vesicular exocytosis with subpixel precision^16^. We applied this approach to directly compare locations of synchronous and asynchronous release events within the same presynaptic bouton. By using quantal analysis, we first selected single-vesicle fusion events (Fig. 3A). We next generated the corresponding ‘Event images’ for sub-pixel localisation of release sites, by applying pixel-by-pixel temporal deconvolution of the unitary SF-iGluSnFR response to the original image time series (Fig. 3B and Methods). The deconvolution procedure allowed us to specifically extract and amplify the spatial component SF-iGluSnFR fluorescence signal associated with the vesicular exocytosis event. This increased the signal-to-noise ratio and enabled us to localise positions of individual release events with approximately 75 nm precision (50 – 100 nm range, Fig. 3C, Fig. S6 and Movie S2). In 95% of boutons, vesicular exocytosis sites were clustered within a single compact area (*S* = 0.09±0.01 μm^2^ in wild type and *S* = 0.10±0.01 μm^2^ at Syt1 KO synapses, Fig. 3D,E and Fig. S7), whilst the remaining 5% of boutons contained two separate release areas. We note that due to exclusion of multivesicular release events and single vesicle events that could not be localised with sufficient precision (*i.e*. within the specified 100 nm threshold, Methods and Fig. S6) we could only determine locations of approximately 50% of all events. Furthermore, the imaged boutons were randomly tilted with respect to the microscope axes. Therefore the release area determined with SF-iGluSnFR on the 2D image projection represents a lower limit of the corresponding active zone area^21^. Indeed, the average active zone size determined using 3D cryo-electron microscopy was ~ 1.35-fold larger (~ 0.12 μm^2^) than obtained with our approach. The release area size varied widely among presynaptic boutons and, as expected, correlated with the total number of vesicles released per action potential (*n_T_*) both in wild type and in Syt1^-/-^ synapses (Fig. 3E).

**Fig. 3.**
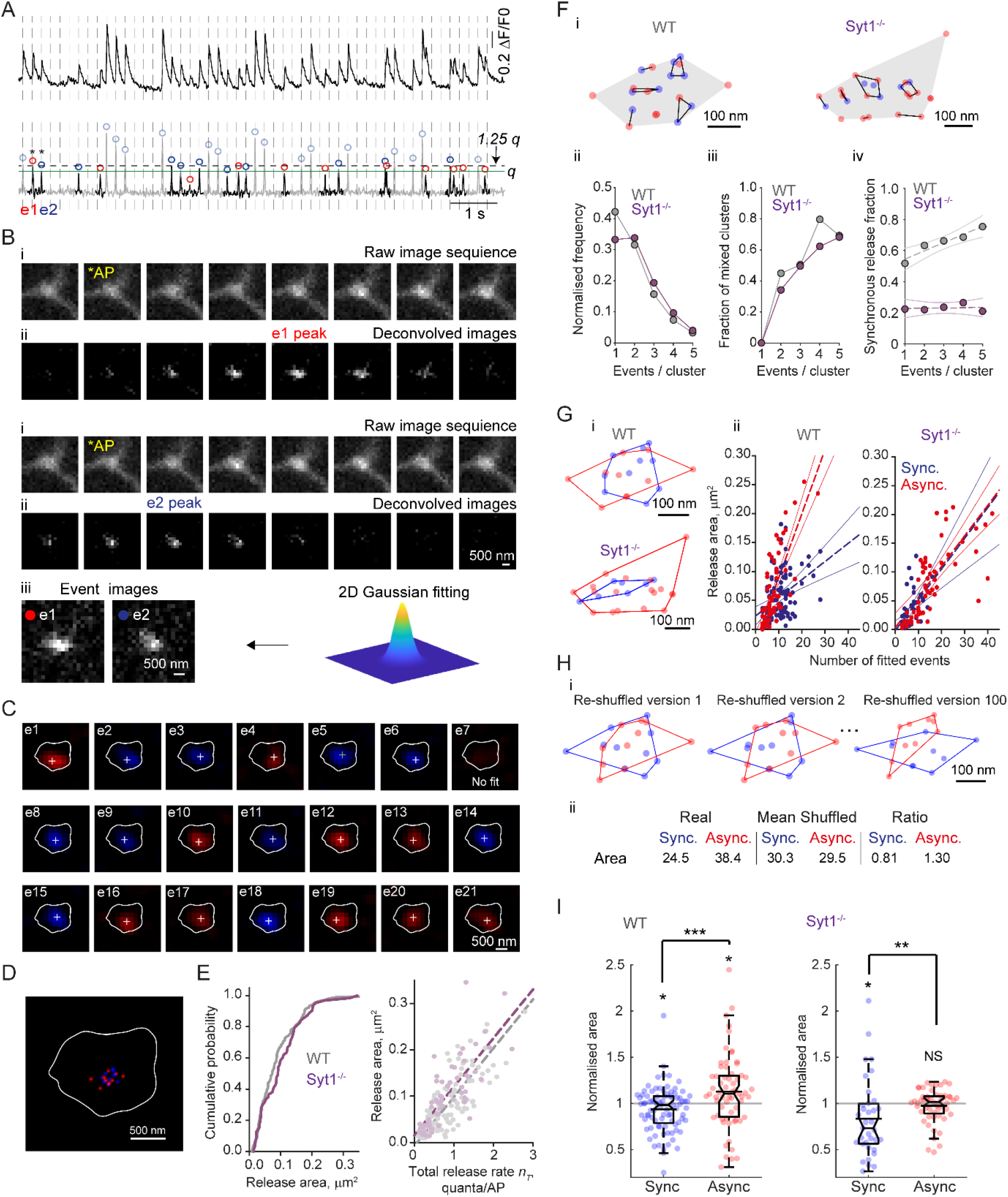
Spatial distribution of synchronous and asynchronous release events in the active zone. **(A** to **D)** Illustration of sub-pixel localisation analysis for single vesicle exocytosis events. (A) Raw (top) and deconvolved (bottom) SF-iGluSnFR traces from a representative bouton. Only single vesicle release events (black areas on the deconvolved trace) were included into the localisation analysis. These were selected using 1.25 *q* amplitude threshold (dashed line), where *q* (green line) is the quantal size calculated as detailed in Fig. 1E. (B) Image pre-possessing in time domain. (i) Raw image sequences before and after event 1 (e1, asynchronous release, 3 frames after the preceding action potential, *AP) and event 2 (e2, synchronous release, 1 frame after the preceding action potential). (ii) Corresponding image sequences after pixel-by-pixel deconvolution of the unitary SF-iGluSnFR response. (iii) ‘Event images’, obtained by averaging 3 frames centred at the response peak on the deconvolved image stack used for sub-pixel localisation of release locations using 2D Gaussian fitting (Methods). (C) Sub-pixel localisation (white crosses) of single quantal events selected in (A) (see also Movie S2). Images are after application of a wavelet filter (Methods); e7 could not be localised due to a low signal-to-noise ratio. The bouton outline (white line) was determined by calculating maximal projection of the deconvolved image stack. (D) Composite image showing relative locations of synchronous (blue) and asynchronous (red) release events within a compact release area. (**E**) Distribution of release area sizes and their dependency on the overall release rate (n = 106 boutons from N=16 cells for wild type and n = 64 boutons from N=11 cells for Syt1^-/-^ neurons). (**F**) Hierarchical cluster analysis of vesicular exocytosis sites. (i) Distributions of clustered events (clustering diameter threshold 100 nm, see Methods) in representative wild type (same as in A-D) and SytT^-/-^ boutons. (ii) Distribution of cluster sizes (pooled data from m = 799 clusters from n = 106 boutons in wild type and m = 553 clusters from n = 64 boutons in Syt1^-/-^ neurons). (iii) Dependency of mixed clusters fraction (containing both synchronous and asynchronous events) on the cluster size. (iv) Increase of synchronous events fraction with cluster size in wild type synapses. Dashed lines, linear regression. Solid lines 95% confidence intervals. (**G**) Comparison of convex hull areas circumventing synchronous or asynchronous events. (i) Representative boutons (as shown in F). (ii) Relationship between the release areas and the number of fitted events in all analysed boutons. Dashed lines, linear regression. Solid lines 95% confidence intervals. (**H**) Illustration of the reshuffling analysis used to compare spatial distributions of synchronous and asynchronous events within individual synapses (representative re-shuffled versions of the wild type bouton shown in A-G, see text for details). (**I**) Distribution of the normalised areas for synchronous and asynchronous events in wild type and Syt1^-/-^ synapses. * p < 0.05, *** p < 0.001, NS p > 0.2, Mann–Whitney U test.

To compare the relative locations of synchronous and asynchronous exocytosis sites, we applied hierarchical cluster analysis (Fig. 3F and Methods). In line with previous studies^2, 21^, locations of individual release events could be grouped into small clusters (<100 nm diameter), which likely correspond to distinct release sites that are reused during repetitive stimulation. We found that among clusters that had at least two events, ~49% of clusters in wild type and ~43% of clusters in Syt1^-/-^ neurons contained both synchronous and asynchronous events. This argues that locations of synchronous and asynchronous events overlap and that they can occur from the same sites. However, in wild type neurons, the fraction of synchronous events progressively increased with the number of events in the cluster. Furthermore, the area covered by synchronous events had a shallower dependency on the number of events (slope 0.003 μm^2^ / event) than the area covered by asynchronous events (slope 0.011 μm^2^ / event). Together these findings suggest that synchronous release is confined to narrower nanodomains within the active zone. To test this at the level of individual synapses we compared each bouton against reshuffled versions of itself (Fig. 3G, H). In each simulation we maintained the number of synchronous and asynchronous events detected in each bouton but randomly scrambled their locations. We next calculated the average areas covered by each release mode (among all simulations) and used these to normalise the experimental values. Our rational was as follows: normalised value greater than 1 indicates that real events have a sparser distribution than the random counterpart, whilst normalised value smaller than 1 indicates that the real events have a more compact distribution. The re-shuffling analysis showed that the normalised area was indeed greater for asynchronous than for synchronous release (Fig. 3H, I). We therefore conclude that asynchronous events in wild type synapses are more widely dispersed across the release area than synchronous events. In contrast, the overall prevalence of asynchronous release in Syt1^-/-^ neurons, largely occluded the difference between spatial distributions of synchronous and asynchronous exocytosis sites, albeit more compact clustering of the remaining synchronous release events was still detected with the re-shuffling analysis (Fig. 3I).

Which mechanisms could explain the differential regulation of synchronous and asynchronous exocytosis at inter- and intrasynaptic levels? Our finding that locations of synchronous and asynchronous events overlap to a large extent, argues that both release modes can occur from the same pool of vesicles. On the other hand, a wider distribution of asynchronous release loci indicates that the probability that a given vesicle will be released synchronously or asynchronously depends on its location within the active zone.

A simple model that can potentially explain our findings, is that the observed variability of release properties could be a consequence of heterogeneity in coupling distances between VGCC and RRP vesicles. The synapse-specific tuning of the coupling distance has been demonstrated in synaptic outputs of pyramidal cells in the neocortex and has been implicated in the target cell-specific regulation of release probability and short-term plasticity^22^. The probability of synchronous release steeply depends on the coupling distance between release-ready vesicles and VGCCs, because it is triggered by transient Ca^2+^-nano/microdomains with a steep spatial gradient. In contrast, asynchronous release is not expected to depend on the coupling distance, as it is triggered by a more global (‘residual’) increase in the presynaptic Ca^2+^ concentration (Fig. 4A)^1^. Therefore, it follows that an increase of coupling distance should lead (i) to a decrease of the overall release probability with concurrent increase of asynchronous release fraction and (ii) to a wider distribution of vesicular exocytosis within the active zone.

**Fig. 4.**
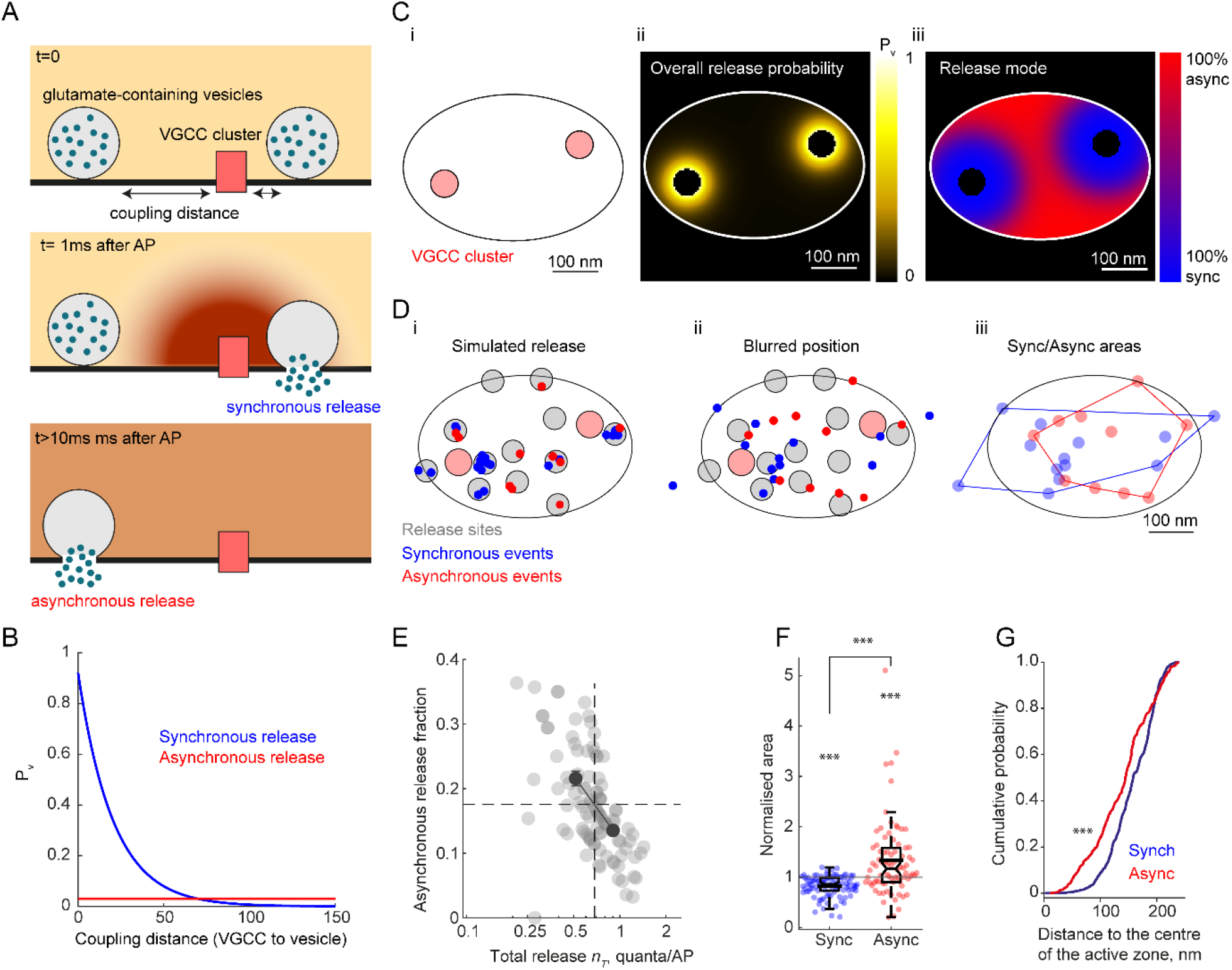
Modelling of synchronous and asynchronous release – implementation of the ‘variable coupling distance’ model. (**A**) Ca^2+^ triggering of different release modes. Synchronous release occurs within several milliseconds after an action potential and is triggered by transient local Ca^2+^- nano/microdomains (~10-100 μM range) in the close vicinity of activated VGCCs (20 – 100 nm range). In contrast, asynchronous release occurs on a time scale of tens to hundreds of milliseconds and is triggered by longer-lasting global changes in presynaptic Ca^2+^ concentration (~ 1-5 μM range)^1^. (**B**) Model predicted dependencies of synchronous (blue trace) and asynchronous (red trace) release probabilities on the distance to a VGCC cluster (see Methods). (**C**) (i) Typical ellipse-shaped active zone containing two VGCC clusters, (ii) model-predicted maps of the overall release probability, and (iii) of the relative fractions of synchronous and asynchronous release modes. (**D**) Results of a representative simulation. (i) Grey circles depict randomly assigned locations of vesicular release sites. Blue and red dots correspond to synchronous and asynchronous release events respectively, which occurred during a simulated train of 51 action potentials. (ii) To mimic the experimental data, locations of release events were blurred by adding a random value corresponding to the accuracy of SF-iGluSnFR event localisation (75 nm). (iii) Comparison of convex hull areas circumventing simulated locations of synchronous or asynchronous events. (**E, F**) Comparison of the model’s output to the experimental results. The simulated data from n = 100 runs (boutons) were processed using the same analysis routine as for the experimental data set. (E) Model-predicted relationship between asynchronous release fraction and the overall vesicular release rate *n_T_* (similar to analysis in Fig. 2A). (F) Results of reshuffling analysis (similar to Fig. 3H, I). (**G**) Distribution of distances to the centre of the active zone for simulated synchronous and asynchronous release events. In agreement with recent electron microscopy data^4, 12^ the model predicts that locations of asynchronous release events are expected to be biased towards the centre of the active zone. * p < 0.05, *** p < 0.001, NS p > 0.2, Mann–Whitney U test.

To test whether our results are consistent with this hypothesis we performed experimentally constrained modelling of presynaptic Ca^2+^ dynamics and activation of vesicular fusion (Fig. 4B-G and Methods). The model was based on the recent functional electron microscopy and single-molecule localisation data showing that presynaptic VGCCs, synchronous release sites and postsynaptic AMPA receptors colocalise within 20 – 30 nm distance and that AMPA receptor clusters preferentially localise to the periphery of the postsynaptic density/active zone^2^’^4, 12^. The model reproduced both a greater asynchronous release fraction at low release probability boutons (Fig. 4E) and a wider distribution of asynchronous release loci within the release area (Fig. 4F), thus arguing that synapse specific adjustment of coupling distances between VGCCs and vesicular release sites could be a major factor that accounts for the differential regulation of synchronous and asynchronous release.

At first sight, our findings contradict the electron microscopy studies, which demonstrated that asynchronous release loci are more biased towards the centre of the active zone^4, 12^. Synchronous release events are expected to concentrate around VGCC clusters, which are predominantly located at the periphery of the active zone^2–4^ (Fig. 4C). Then it follows that asynchronous events, which are located further away from the VGCC clusters, can indeed be positioned closer to the active zone centre but be more widely distributed across the release area than synchronous events (as supported by our results).

Accumulating data demonstrate that the coupling distance between VGCCs and release-ready vesicles is dynamically regulated by presynaptic scaffold proteins (in particular RIM and RBP2)^23, 24^ and it remains to be established whether and how this signalling pathway is regulated by the identity of the postsynaptic target cell. It also remains to be determined whether the balance between different release modes in synapses supplied by the same axon is further tuned by the adjustment of the local abundance of different synaptotagmin isoforms. In summary, our results reveal a general principle that relates the two major functional properties of small central glutamatergic synapses – the release probability and the synchronicity of vesicular release. We demonstrate that the ratio between synchronous and asynchronous synaptic vesicle exocytosis varies extensively among presynaptic boutons supplied by the same axon, and that asynchronous release fraction is elevated in parallel with short-term facilitation at synapses with low release probability. Such fine-tuning of different release modes, together with target cell-specific adjustment of release probability and shortterm plasticity^7, 22, 25, 26^, has the potential to provide vast flexibility for synaptic computations in neural circuits composed of different cell types.

## Materials and Methods

### Neuronal cultures and SF-iGluSnFR expression

Experiments conformed to the Animals (Scientific Procedures) Act 1986, and were approved by the UK Home Office. Primary cortical neurons were produced from either wild type (C57BL/6J; Charles River) or Syt1^-/-^ (B6; 129S-Syt1tm1Sud/J; The Jackson Laboratory) postnatal day 0 mouse pups of both sexes and cultured in Neurobasal A/B27-based medium (Thermo Fisher Scientific). The cortices were dissected and dissociated by enzymatic digestion in 0.25% trypsin for 10 min at 37°C and then triturated using a standard p1000 micropipette. Neurons were plated on poly-L-lysine-treated 19-mm glass coverslips (1 mg/mL; Sigma-Aldrich) at a density of ~100,000 cells per coverslip. At 5 days in vitro (5 DIV) neurons were transfected with pAAV.hSynap.SF-iGluSnFR.A184V plasmid ^14^ (addgene Plasmid #106174) using Neuromag reagent (KC30800; OZ Biosciences). The transfection resulted in sparse expression of the iGluSnFR probe in a small subpopulation of neurons (~3%), which allowed us to select individual cells for imaging. Experiments were performed between 16 and 21 DIV.

### SF-iGluSnFR imaging of glutamate release

#### Experimental set up

SF-iGluSnFR fluorescence imaging experiments were performed on an inverted Olympus IX71 microscope equipped with a Prime95B back illuminated CMOS camera (Teledyne Photometrics) using a 60x oil-immersion objective (1.35 NA) with resulting image pixel size 183.3 nm. SF-iGluSnFR fluorescence was recorded using a 470-nm excitation light-emitting diode (OptoLED Light Source, Cairn Research) and a 500–550 band-pass emission filter. Image acquisition was performed using μManager software^27^.

Experiments were conducted in a custom-made open laminar flow perfusion chamber (volume 0.35 ml, perfusion rate ~ 1 ml/min) at 23–25 °C. The imaging extracellular solution contained (in mM): 125 NaCl, 26 NHCO_3_, 12 Glucose, 1.25 NaH_2_PO_4_, 2.5 KCl, 2 CaCl_2_, 1.3 MgCl_2_ (bubbled with 95% O2 and 5% CO_2_, pH 7.4). To ensure that recorded SF-iGluSnFR responses originate only from the stimulated axon, we suppressed recurrent activity in the neuronal network by blocking postsynaptic ionotropic glutamate and GABA receptors with (in μM) 50 DL-AP5 (Abcam), 10 NBQX (Abcam), and 50 Picrotoxin (Tocris Bioscience).

#### Electrophysiology

A putative pyramidal-like neuron expressing the SF-iGluSnFR probe that did not contain any other transfected cells in its vicinity was selected for imaging. A whole-cell voltage clamp recording was established in the selected cell using a fire-polished borosilicate pipette (4-7 MΩ, Warner Instruments) and Axon Multiclamp 700B amplifier (Molecular Devices). The intracellular pipette solution contained (in mM): 105 K^+^ Gluconate, 30 KCl, 10 HEPES, 10 Phosphocreatine-Na_2_, 4 ATP-Mg, 0.3 GTP-NaH_2_0, 1 EGTA (pH=7.3, balanced with KOH). The Multiclamp commander software was used to measure the series resistance (~20 MΩ), which was compensated at ~30%. Signal was acquired at 20 kHz (4-kHz Bessel-filtering) using a National Instrument board NI USB-6221 controlled with WinWCP software (created and provided by John Dempster, University of Strathclyde). A liquid junction potential of −10mV was subtracted from the measurements *post hoc*. The recorded neuron was held at −70mV and action potentials were evoked using brief voltage steps (5 ms) from −70mV to −10mV, producing stereotypical ‘escape’ currents.

#### Fluorescence imaging

After establishing a patch-clamp recording, evoked synaptic SF-iGluSnFR responses were imaged in 259 x 23μm (1412 x 125 pixels) ROIs located within the axonal arbour 200 – 1000 μm away from the soma. Images were acquired at 250 Hz sampling rate (4 ms/frame). The exposure time of each frame was captured with the data acquisition board. This allowed us to align the timing of captured frames to the electrophysiological recording, and therefore to determine the time between each vesicular release event and the preceding to action potential. For each neuron between 1 to 4 different ROIs were imaged with approximately 5 minutes interval between the trials. The stimulation protocols consisted either of 51 action potentials delivered at 5 Hz, or of 10 pairs of action potentials delivered at 20 Hz (50 ms interspike interval) with 10 seconds between individual paired pulse sweeps.

### Image analysis

Image analysis was performed offline using ImageJ (NIH)^28^ and MATLAB (MathWorks) custom-developed scripts.

#### Identification of active boutons

In order to detect all active presynaptic boutons located within the chosen ROI, a series of filters was applied to the acquired image stack in the X-Y (space) and Z (time) dimensions. First, for each pixel, a moving average filter with a 3-point span was used to smooth the temporal profile of the SF-iGluSnFR responses. Next, a bandpass Gaussian filter (0.5 Hz – 30 Hz) was applied in order to amplify the SF-iGluSnFR signal, revealing glutamate release sites by removing background fluorescence (Fig. 1C, D and Movie S1). Next, a median filter (3×3 pixels) was used to reduce the spatial high-frequency noise component and to improve the robustness of automatic detection of active boutons. Finally, a maximal projection of the filtered stack was obtained in order to visualise all regions where glutamate release events were detected, regardless of the rate of their occurrence. Positions of putative glutamate release sites on the maximal projection image were automatically detected using Find Maxima ImageJ plugin and a set of circular ROIs centred on the detected maxima (diameter 5 pixels, ~0.9μm) was created. The detection threshold was chosen in a such way that all putative release sites were included, along with false-positive sites originating from background noise (these were excluded during quantal analysis stage as described below).

#### Detection of quantal glutamate release events

To determine the timings and the amplitudes of vesicular release events the SF-iGluSnFR fluorescence signal from each selected ROI was first filtered using a 0.5 Hz – 30 Hz bandpass Gaussian filter, which allowed us to remove the baseline drift and the high frequency noise. This was followed by deconvolution of the experimentally determined average unitary SF-iGluSnFR response, approximated as an instantaneous rise followed by an exponential decay 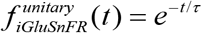 with a decay time *τ* = 68 ms (Fig. S1): 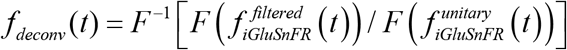, where *F* is the discrete Fourier transform and *F*^-1^ is the inverse Fourier transform functions (MATLAB). The obtained deconvolved trace was further filtered using a 0.5 Hz – 30 Hz bandpass Gaussian filter to improve signal-to-noise ratio. In line with previous reports^17, 29^, the all-point histogram of the deconvolved trace could be well-approximated by a single Gaussian centred at zero: 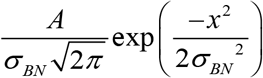 (Fig. S2A). The obtained standard deviation *σ_BN_* (characterising the level of baseline noise) was then used to set the bouton-specific threshold for the detection of quantal events at *θ* = 4*σ_BN_* ^29^. The amplitudes and the timings of individual release events were then determined at local maxima on the filtered deconvolved trace above the threshold.

#### Quantal analysis

To determine the amplitude of deconvolved SF-iGluSnFR signal corresponding to release of a single quanta (*q*) and the distribution of quantal vesicular release events in each bouton, we generated a quasi-continuous amplitude histogram using a bootstrapping procedure with added bouton specific noise (*m* = 100,000 simulations). The simulated values were obtained by randomly selecting an experimentally determined amplitude and adding a random number form a Gaussian distribution with a standard deviation 2*σ_BN_*. The positions of quantal peaks on the obtained histogram were fitted using a finite mixture model consisting of the sum of Gaussians: 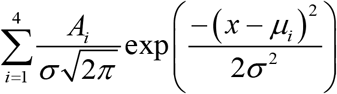, where *A_i_* is the amplitude of the *i*th peak, 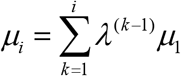 is the average SF-iGluSnFR event amplitude corresponding to simultaneous release of *i* vesicles, *λ* is a factor that accounts for possible progressive saturation of SF-iGluSnFR signals during multi-vesicular release events, *σ*^2^ = *σ_BN_*^2^ + *σ_A_*^2^ is the variance of each peak, and *σ_AN_*^2^ is the noise component associated with variability of SF-iGluSnFR amplitudes (including the variability caused by a random jitter between the timings of release events and the camera exposure cycle, Fig. S3). The use of bootstrapping allowed us to eliminate the error associated with the sensitivity of fitting procedure to the histogram bin size. The mean value of the saturation factor *λ* estimated across n = 1,130 boutons from 16 WT neurons was 0.9 (coefficient of variation 0.15). Thus, in line with a previous study in ribbon synapses^17^, the amplitudes of deconvolved SF-iGluSnFR signals provided a nearly linear read-out for multi-vesicular release.

#### Temporal resolution and sensitivity of quantal analysis

To estimate the temporal resolution and the sensitivity of SF-iGluSnFR quantal analysis we simulated a set of SF-iGluSnFR synaptic responses (n = 2,000) using the experimentally determined signal-to-noise ratio (Fig. S2B) and processed the obtained traces using the filtering and deconvolution procedures described above. Analysis of simulated traces verified that temporal resolution of detection of vesicular release events was primarily limited by the camera acquisition rate (4 ms). The simulations confirmed that the use of *θ* = 4*σ_BN_* detection threshold effectively abolished the presence of false-positive events (5 false positive in 1000 simulated traces). However, the fraction of false-negative (missed) events was increased with the decrease of signal-to-noise ratio (Fig. S3E). We therefore excluded boutons with signal-to-noise ratio below 5*σ_BN_* (~ 17% of all boutons, which are likely to be boutons outside the focal plane), thus limiting the fraction of false-negative events to 20%.

#### Sub-pixel localisation of vesicular release sites

For each active bouton a 21 x 21-pixel ROI (3.85 x 3.85 μm), centred at the corresponding intensity maximum on the maximal projection filtered stack (Fig. 1D), was extracted from the raw SF-iGluSnFR image stack. If the distance between two neighbouring boutons was less than 16 pixels (~ 3 μm) they were excluded from the analysis.

To increase the signal-to-noise ratio, the extracted image stacks were pre-processed pixel-by-pixel in time domain. The traces were processed using a bandpass Gaussian filter (0.5 Hz – 30 Hz), which was followed by deconvolution of the unitary SF-iGluSnFR response from the filtered signal, as described in the ‘Identification of active boutons’ section. Vesicular release event was considered to contain a single quantum if its amplitude was below a 1.25 *q* threshold (determined using quantal analysis). For each single quantal release, an ‘event image’ was calculated by averaging 3 frames from the deconvolved image stack centred at the response peak (Fig. 3B). The event image was then used for sub-pixel localisation of the vesicular release site with the ThunderSTORM ImageJ plugin^30^.

The analysis consisted of two steps: finding the approximate position of the release site and its subsequent sub-pixel localisation. The event image was filtered using a wavelet filter, with a B-spline function of order 3 and a scale of 2. The approximate position of the release site was determined by finding a local maximum on the filtered image (intensity is greater than the specified threshold and at the same time greater than or equal to the intensities of 8 neighbouring pixels). Because the signal-to-noise ratio varied among synapses, the optimal threshold value was set independently for each bouton. For this purpose, we generated a ‘background noise’ image stack consisted of 1,000 images. Each ‘background noise’ image was obtained by averaging 3 randomly selected frames from the deconvolved image stack, which were separated from the nearest release event by at least 15 frames (or 60 ms). The optimal bouton-specific threshold was then determined by running the analysis iteratively on the ‘background noise’ image stack, starting from a high level of intensity and decreasing it by half at every iteration until some events were detected. A second iterative process increased the threshold value in 250 intensity level steps, until no maxima were detected. The determined threshold value was next used to analyse the event images from the same bouton. Event images where the approximate position of the release site could not be determined with the selected threshold were excluded from the analysis. Because we preselected images corresponding to release of single vesicles, we *a priori* knew that there should be only one spatial maximum in each image, therefore we excluded images if more than one local maximum was detected. Finally, the sub-pixel localisation of single quanta release events that passed all the above selection criteria was performed by fitting an integrated form of 2D Gaussian function into the 7 x 7 pixel array centred around the established approximate positions^30^.

#### Estimation of release sites localisation precision

The precision of localisation of vesicular release sites depends on several key factors. First as in the case of single molecule localisation microscopy, the localisation accuracy depends on the spatial resolution (point spread function, PSF) of the microscope and on the strength of SF-iGluSnFR signal (*i.e*. the signal-to-noise ratio determined by the number of photons collected for a given release event). In addition to this, the localisation accuracy of release events also depends on the spatial profile of the activated SF-iGluSnFR molecules within a given active zone and on the relative position and tilt of the imaged active zone with respect to the microscope focal plane^21^.

To account for the joint effect of all the above factors we estimated the localisation accuracy using an empirical approach. For each event we generated an ‘added noise’ image stack (in total 50 images). Each of the images in the ‘added noise’ stack was a sum of the original event image and a randomly selected image from the ‘background noise’ image stack described in the ‘Sub-pixel localisation of vesicular release sites’ above. We next applied the localisation analysis to individual images from the ‘added noise’ stack and calculated the average position of all fits (Fig. S6B). The distance *δ* between the initial fit and the average position obtained from the ‘added noise’ image stack was used as an empirical estimate of the localisation accuracy (Fig. S6 C, D) and events with *δ* above the 100 nm threshold were excluded from analysis.

To test if *δ* represents a realistic estimate of the localisation accuracy we performed similar analysis using artificial computer-simulated images. The rational was that in this case we *a priory* knew the true locations of release sites. The simulated ‘event images’ were obtained using the following steps. We first simulated a spatio-temporal SF-iGluSnFR profiles corresponding to single vesicle fusion events using a general function form:

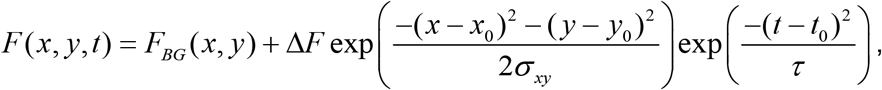

where (*x*_0_,*y*_0_) and *t*_0_ are coordinates and timing of a release event; *τ* = 68 ms, SF-iGluSnFR decay rate (Fig. S1), *σ_xy_*, spatial width of SF-iGluSnFR response (randomly selected from the experimentally determined range 250-450 nm), *F_BG_* (*x*, *y*) and Δ*F* background SF-iGluSnFR fluorescence and the amplitude of responses respectively (randomly selected from the recoded boutons). The obtained *F*(*x*,*y*,*t*) responses were mapped on the experimental spatio-temporal grid (183.3 nm in space and 4 ms in time domain) and noise was added using a randomly selected value of signal-to-noise ratio from the experimental data set (Fig. S2B). Finally, locations of simulated vesicular release events were fitted using the same routine as for the experimental data. The analysis of artificial images verified that *δ* provides an accurate estimate of localisation precision (50 – 100 nm range). Furthermore, the computer simulations demonstrated that the use of the averaged position determined from the ‘added noise’ stack provides a more reliable estimate for the location of release site. Therefore, we used this improved averaged fit in the analysis of the relative distributions of synchronous and asynchronous release events in Fig. 3F-G.

#### Hierarchical cluster analysis

Hierarchical cluster analysis was used to investigate the distribution of synchronous and asynchronous events among different release sites (Fig. 3F and Fig. S7). The built-in MATLAB functions `‘linkage’` and `‘cluster’` were used to best group events based on a distance threshold, allowing individual release cites to be defined by events located within 100 nm of the cluster centroid (in accordance with the precision of localisation of individual exocytosis events). The same algorithm was applied to identify synapses with two active zones (see examples in Fig S7). If boutons displayed event clusters with centroids distanced by 700 nm or more, they were defined as having two active zones and were excluded from spatial analysis.

### Computational modelling of synchronous and asynchronous release within the active zone

Based on the available electron microscopy data^3, 4, 12, 31, 32^, we considered a typical active zone of an area *S* = 0.12 μm^2^, that contained two VGCCs clusters (located at the active zone periphery) and *N_rel_* = 12 of vesicular release cites that were randomly distributed across the active zone (Fig. 4C,D). The modelling of three-dimensional action potential-evoked presynaptic Ca^2+^ dynamics and of Ca^2+^ activation of vesicular release was performed using the presynaptic terminal model implemented in the Virtual Cell (VCell) simulation environment (http://vcell.org) and custom-developed MATLAB (MathWorks) scripts as described in detail in our previous work^33–35^. Briefly, we simulated action potential-evoked spatio-temporal Ca^2+^ dynamics in the vicinity of a VGCC cluster (containing 7 P/Q-type, 8 N-type, and 1 R-type VGCCs) in the presence of the three endogenous buffers (calmodulin, Calbindin D28k and ATP). We next computed the probability maps for synchronous release in response to Ca^2+^ influx at each of the two VGCC clusters (*p*1*_v_* (*x*, *y*) and *p*2*_v_* (*x*, *y*)) using the six-state allosteric model of Ca^2+^ activation of vesicle fusion^36^. As there was virtually no spatial overlap between *p*1*_S_*(*x*, *y*) and *p*2*_S_*(*x*, *y*) we approximated the synchronous release probability map as *p_s_*(*x*, *y*) = *p*1_*S*_(*x*, *y*) + *p*2_*s*_(*x*, *y*). Synchronous release was then simulated by considering the overall release probability as the product *p_S_*(*x*, *y*) · *p_oc_*, where *p_oc_* is the probability that a given release site is occupied by a RRP vesicle. Because vesicular release during 5 Hz stimulation was on average depressed by ~ 2 fold in comparison to the response at the 1^st^ action potential, we considered *p_oc_* =0.5. Probability of asynchronous release was assumed to be independent on the release site location and was estimated as *p_A_* = *n_A_*/*N_rel_* = 0.01, where *n_A_* is the average rate of asynchronous release per action potential during 5 Hz stimulation (*n_A_* ~ 0.12, Fig. 1G).

### Data Inclusion and Exclusion Criteria

In addition to the exclusion criteria described in the above sections we applied the following selection criteria. In Fig. 2A, B, boutons with at least 2 release events were selected to allow meaningful estimation of *n_A_*/*n_T_*. Similarly, in Fig. 2C boutons with at least 2 synchronous events were selected to allow estimation of *PPR*. Previous work^21^, where vGlut1-pHluorin probe was used for sub-pixel localisation of vesicular release sites, demonstrated that positions of release events can be reliably fitted only within ~ 100 nm from the imaging plane in z-direction and that selection of boutons with at least 5 detected events effectively limits the analysis to a sub-population of active zones that are mostly parallel to the image plane (within a 20° tilt). We therefore applied the same selection criteria in our analysis in Fig.3. Furthermore, in order to compare the relative locations of synchronous and asynchronous release events (Fig. 3F-G) only boutons that contained both event types were included.

### Statistical analysis

The distribution of data in each set of experiments was first tested for normality using the Shapiro-Wilk test. The similarity of variances between each group of data was tested using the F test. Normally distributed data were presented as mean ± s.e.m., each plot also contained the individual data points. Student’s t tests for group means or paired t tests were used as indicated. The data sets that failed the normality test were presented using box- and – whisker plots (box 25^th^ – 75^th^ percentiles, whiskers 10^th^ – 90^th^ percentiles), each plot also contained the individual data points and compared using Mann-Whitney U test. The detailed statistical analysis is presented in Supplementary Table 1. No statistical methods were used to pre-determine sample sizes, but our sample sizes were similar to those reported in previous publications that use similar techniques^16, 17, 34^. Data analysis was performed blind to the conditions and genotype tested. All statistical tests were performed using SigmaPlot 11 (Systat Software) and MATLAB (MathWorks) software packages.

## Acknowledgements

We are grateful to Dimitri Kullmann, James Rothman, Shyam Krishnakumar and Christopher Kushmerick for reading the manuscript and providing critical feedback. The study was supported by the Wellcome Trust Strategic Award 104033/z/14/z (K.E.V.), Epilepsy Research UK Project Grant P1806 (K.E.V.), the Wellcome Trust PhD Studentship 203795/Z/16/Z (H.L.), Medical Research Council UK Project Grant MR/T002786/1 (Y.T. and K.E.V.).

## Author contributions

P.R.F.M. and K.E.V. designed experiments; P.R.F.M., E.T., H.L. and D.K. performed experiments; P.R.F.M., C.G.Z.C., Y.T. and K.E.V. developed the data analysis framework and performed the analysis; Y.T. and K.E.V. performed computational modelling; P.R.F.M., Y.T. and K.E.V. wrote the manuscript; Y.T. and K.E.V. obtained funding; K.E.V. managed the project.

## Data and materials availability

All the data are available upon request from the corresponding authors.

## Supplementary Materials

**Fig. S1.**
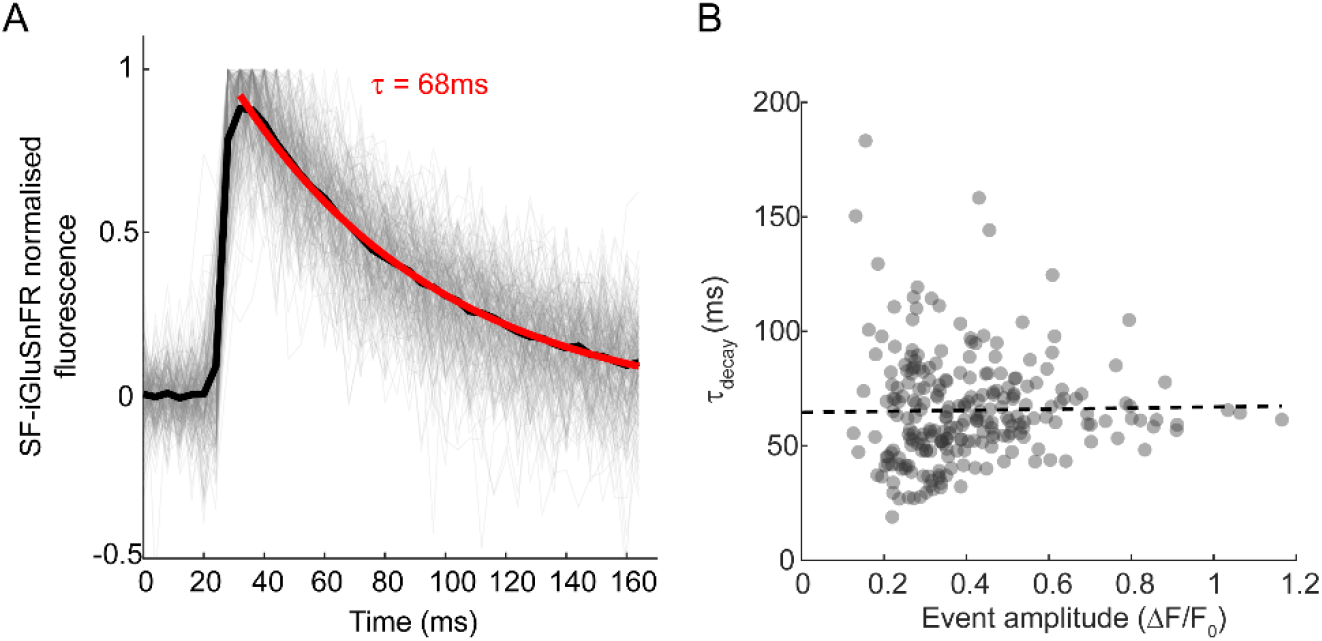
Analysis of kinetics of evoked presynaptic SF-iGluSnFR events. (**A**) Black trace, average of 244 individual action potential-evoked SF-iGluSnFR fluorescence transients (grey traces) aligned by their rising slopes (n = 37 representative presynaptic boutons recorded in N = 6 wild type cells). Red trace, mono-exponential fit of the decay phase of the average trace *F*(*t*) = *e*^-*t/τ*^. (**B**) The values of decay time constant *τ* estimated for individual SF-iGluSnFR events do not correlate with the event amplitude. Dashed line, linear regression.

**Fig. S2.**
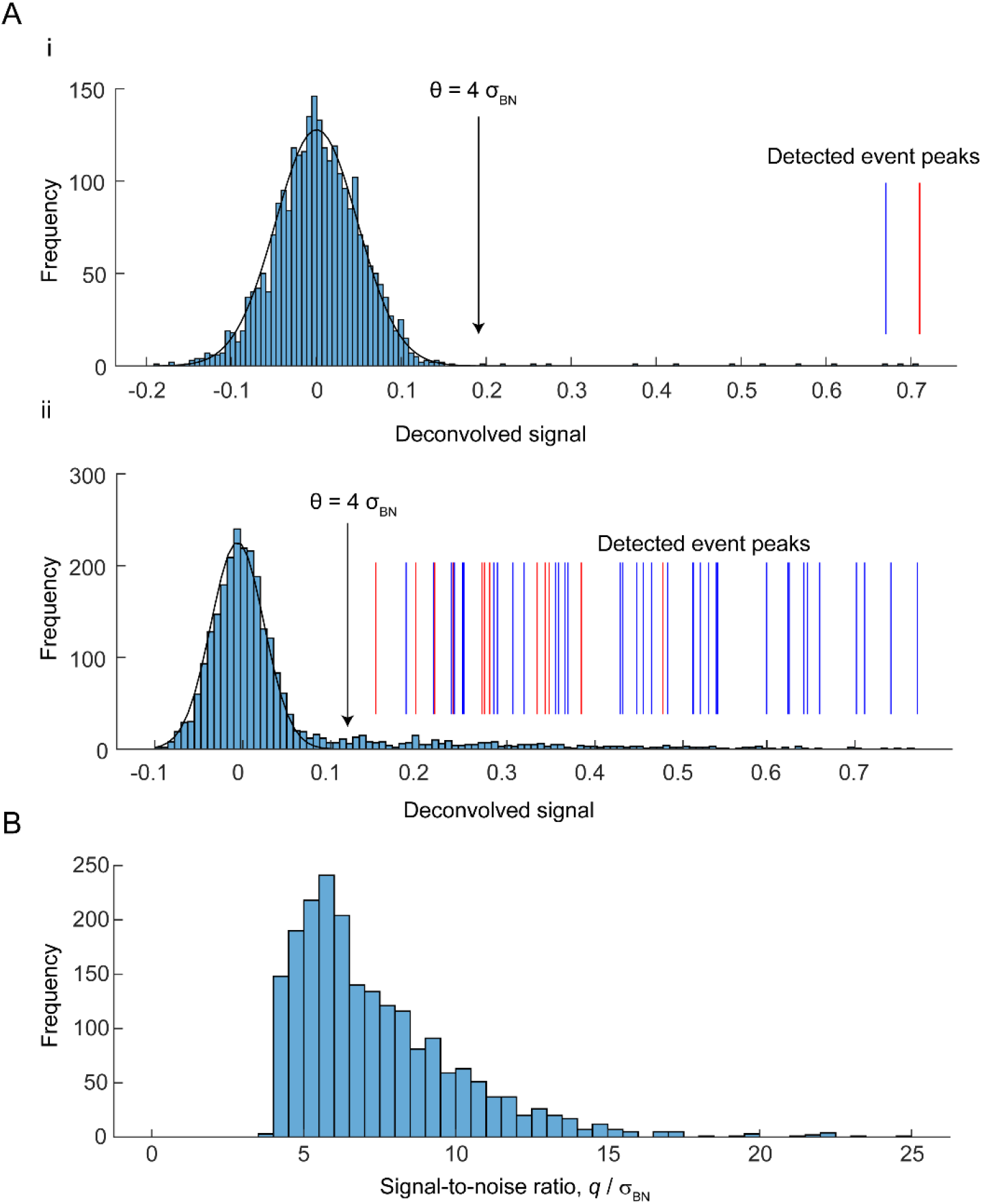
Analysis of background noise and setting a threshold for automatic detection of quantal glutamate release events in single boutons. (**A**) All-point histograms of the deconvolved traces from boutons 1 and 2 in Fig. 1. Black lines, histogram fits with a Gaussian function 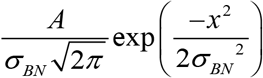. The obtained standard deviation *σ_BN_*, which characterises the baseline noise, was used to set-up the bouton-specific threshold for detection of quantal events: *θ* = 4*σ_BN_* (vertical arrows). Vertical bars depict the amplitudes of SF-iGluSnFR synchronous (blue) and asynchronous (red) release events, determined as local maxima located above the threshold on the deconvolved trace in Fig. 1E. (**B**) Distribution of signal-to-noise ratio across recorded synapses (pooled data from N = 16 wild type and N = 11 Syt1^-/-^ neurons, total n = 2,075 boutons). Signal-to-noise ratio in each bouton was calculated as the ratio of SF-iGluSnFR signal amplitude corresponding to release of a single vesicle (1q) to the standard deviation of the background noise *σ_BN_*.

**Fig. S3.**
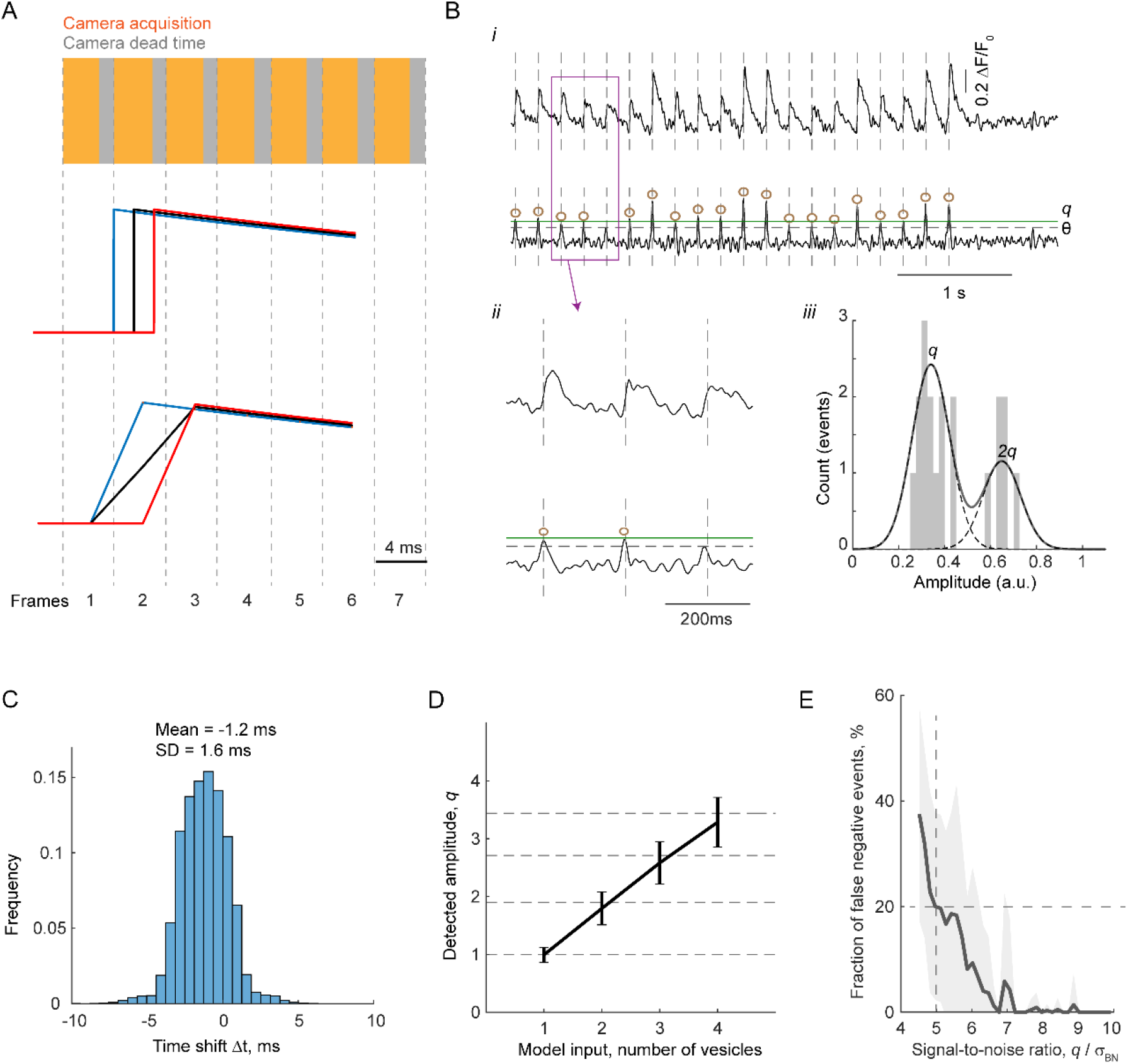
Temporal resolution and sensitivity of quantal analysis. (**A**) Illustration of the effect of a random jitter between the exact vesicular release times and the camera exposure cycle on the profile of SF-iGluSnFR responses. To ensure equal detection of fluorescence signals from all pixels within a selected ROI (typically 1412 x 125) the Prime95B camera was operated in the ‘Pseudo Global Shutter’ mode, when excitation light was only delivered during simultaneous exposure of all selected camera rows. As a result, photons were only collected during the first 2.8 ms (yellow rectangles) of the 4 ms frame cycle (vertical dashed lines). Middle traces illustrate ‘ideal’ SF-iGluSnFR signals that initiate at different times during the frame cycle. Bottom traces depict the shape of corresponding SF-iGluSnFR signals as they would be acquired by the camera. (**B**) Analysis of simulated SF-iGluSnFR responses. Each of 1,000 simulated traces consisted of 1,200 x 4 ms frames. Vesicular release events (20 per trace) were assumed to occur at 5 Hz (*i.e*. every 50^th^ frame). The exact timings of release events within the selected frames were random. The number of vesicles released during each event was drawn from a binomial distribution, assuming *m* = 5 (the number of release ready vesicles) and *p_v_* = 0.15 (release probability of an individual vesicles). The amplitudes of SF-iGluSnFR responses corresponding to simultaneous release of *i* vesicles were approximated as 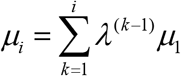, where *λ* = 0.9 is a factor (estimated experimentally) that accounts for a progressive saturation of SF-iGluSnFR during multi-vesicular release (see Methods). The time course of SF-iGluSnFR responses was approximated as an instantaneous rise followed by an exponential decay with a characteristic time *τ* = 68 ms (Fig. S1). The signal-to-noise ratio for each trace was randomly drawn from the experimentally determined distribution (Fig. S2B). (i) Representative band-pass filtered (top) and deconvolved (bottom) simulated SF-iGluSnFR signals with signal-to-noise ratio 5.7 *q*/*σ_BN_*. As in the case of experimental traces (Fig. 1E), release events (brown circles) were identified as local maxima on the deconvolved trace located above the threshold *θ* = 4 *σ_BN_* (horizontal dashed line) corresponding to 4 standard deviations of the background noise. (ii) Enlarged traces corresponding to the boxed area containing a false-negative (missed) event. (iii) Right, quantal analysis of the example artificial trace performed as detailed in Fig. 1E. (**C**) Distribution of the time shifts (△*t*) between the determined and the actual times of simulated events. The results of simulations confirm that temporal resolution of our analysis is limited by the camera acquisition rate (4 ms / frame). The detected event times were systematically shifted to the left by approximately 1.25 ms. This was due to a combined effect of noise, bandpass filtering and the inherent random jitter between the timings of release events and the camera exposure cycle (A). Considering the intrinsic 2 – 6 ms delay in the onset of synchronous release caused by the finite speed of action potential propagation and variable distances between synapses and cell soma^19^ we did not perform correction for this small systematic error. (**D**) Relationship between model input (number of vesicles) and model output (detected amplitude of deconvolved SF-iGluSnFR signal in units of *q*) verifying the robustness of the amplitude histogram fitting approach. Horizontal dashed lines depict model-predicted SF-iGluSnFR amplitude values (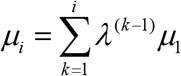, see Methods). (**E**) Dependency of false-negative event fraction on the signal-to-noise ratio. Based on this simulation, boutons with signal-to-noise ratio below 5 *q*/*σ_BN_* (vertical dashed line) were excluded from analysis, thus restricting the fraction of false-negative events to ~20% (horizontal dashed line).

**Fig. S4.**
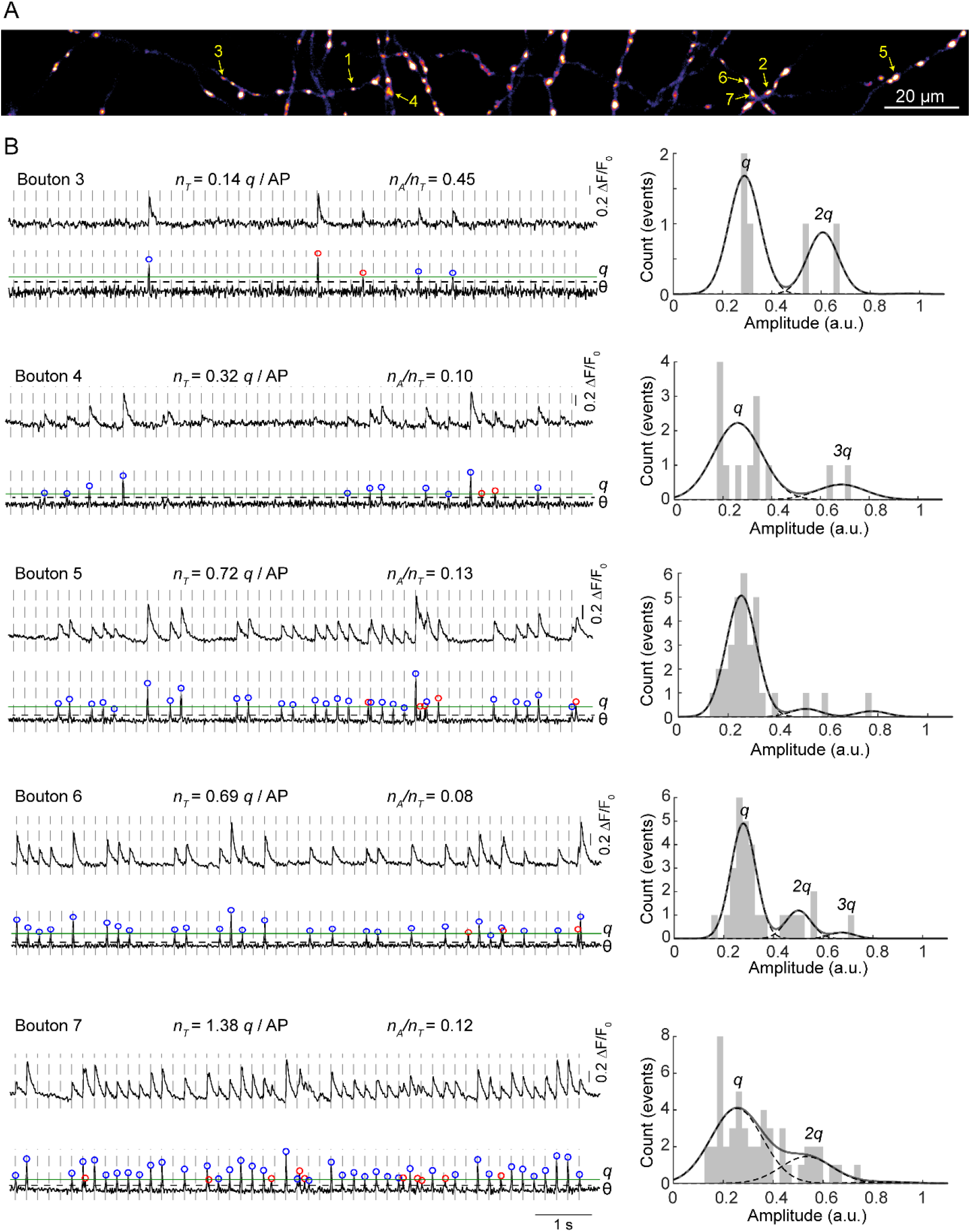
Additional examples of quantal analysis in individual presynaptic boutons (the same wild type neuron as in Fig. 1). (**A**) Heat map of glutamate release sites (maximal projection of the band pass-filtered SF-iGluSnFR image stack, same image as in Fig. 1D). Yellow arrows depict positions of example boutons. (**B**) Quantal analysis of SF-iGluSnFR responses at Boutons 3 to 7 (see Fig.1E for analysis of boutons 1 and 2). Left, band-pass filtered (top traces) and deconvolved (bottom traces) SF-iGluSnFR signals. Vertical dashed lines depict the timings of somatic action potentials. Horizontal black dashed lines – detection thresholds (*θ*) for vesicular release events, corresponding to 4 standard deviations of the background noise. Horizontal green lines – amplitudes of SF-iGluSnFR signals corresponding to release of a single vesicle (single quanta, *q*) determined by fitting positions of peaks on the amplitude histograms (right). Blue and red circles on the deconvolved traces depict synchronous and asynchronous release events respectively. Comparison of quantal responses at boutons 6 and 7 illustrates that SF-iGluSnFR signals in neighbouring boutons are not cross-contaminated by a possible glutamate spillover.

**Fig. S5.**
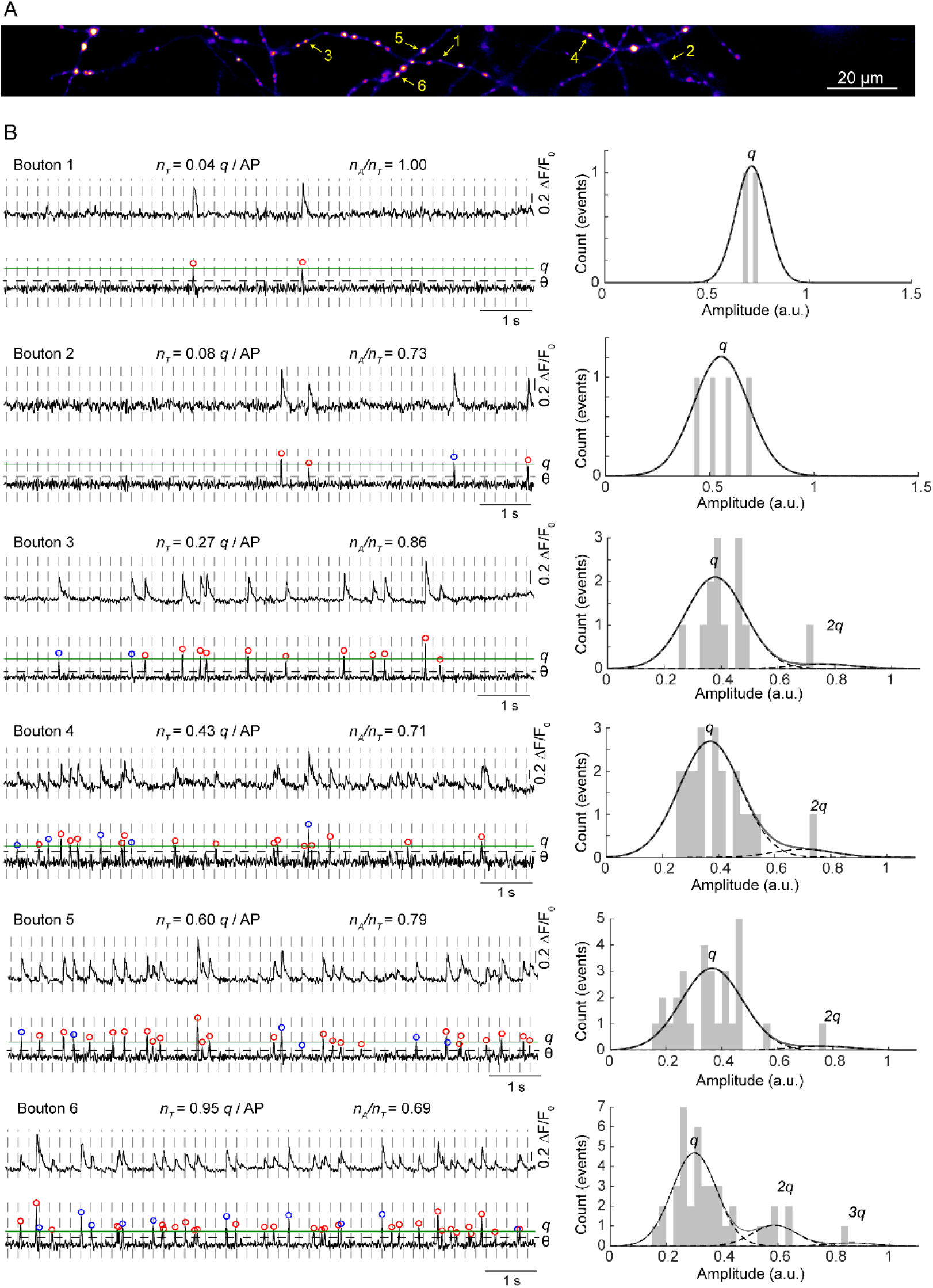
Examples of quantal analysis in individual presynaptic boutons in a representative Syt1^-/-^ neuron. (**A**) Heat map of glutamate release sites (maximal projection of the band pass-filtered SF-iGluSnFR image stack) across the axonal arbour of a Syt1^-/-^ neuron. Yellow arrows depict positions of six example boutons. (**B**) Analysis of SF-iGluSnFR responses in the selected boutons. Left, band-pass filtered (top traces) and deconvolved (bottom traces) SF-iGluSnFR signals. Vertical dashed lines depict the timings of somatic action potentials. Horizontal black dashed lines – detection thresholds (*θ*) for vesicular release events, corresponding to 4 standard deviations of the background noise. Horizontal green lines – amplitudes of SF-iGluSnFR signals corresponding to release of a single vesicle (*q*) determined by fitting positions of peaks on the amplitude histograms (right). Blue and red circles on the deconvolved traces depict synchronous and asynchronous release events respectively.

**Fig. S6.**
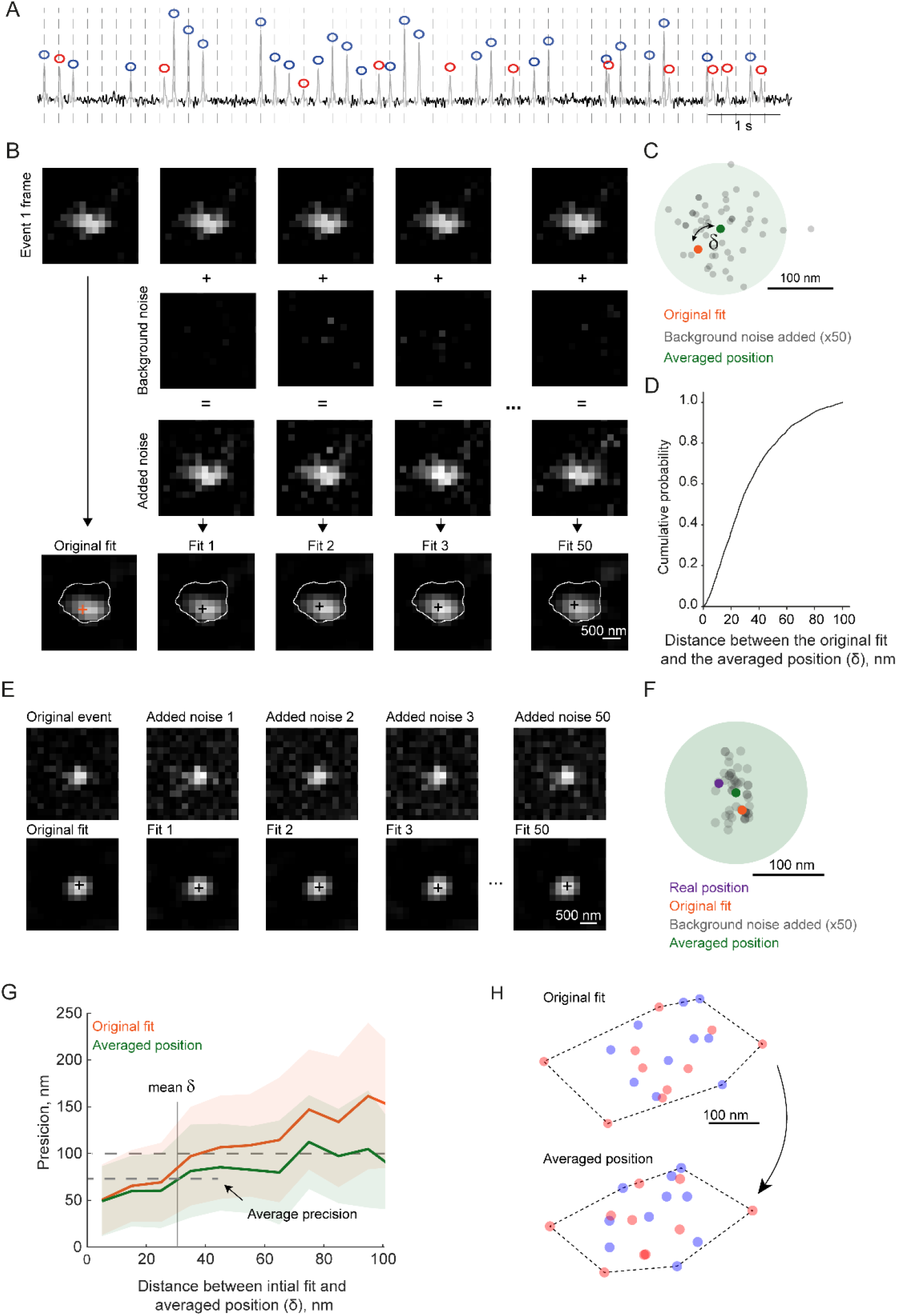
Estimation and improvement of vesicular release sites localisation precision using noise analysis. (**A**) Generation of ‘background noise’ image stack (1000 images). Each background noise image was obtained by averaging 3 randomly selected frames from the deconvolved SF-iGluSnFR image stack, which were separated from the nearest release event by at least 15 frames (depicted in black on the representative trace, the same bouton as in Fig. 3A-D). (**B, C**) Estimation of vesicular release localisation precision (detailed illustration for Event 1 from Fig. 3A-D). (B) For each event, an ‘added noise’ image stack was generated (50x), where each image was a sum of the original ‘event image’ and a randomly selected image from the ‘background noise’ stack. We next run the sub-pixel localisation analysis for each image in the ‘added noise’ stack (grey dots in C), calculated their average position (green dot) and the distance (*δ*) between location of the ‘original’ fit (obtained using the initial ‘event image’, orange dot) and the average position from the ‘added noise’ stack. Events with *δ* above the 100 nm threshold were excluded from analysis. (**D**) Cumulative distribution of *δ* values for 2,592 fitted events from n = 106 wild type boutons, *δ_mean_* = 32.4 ± 23.0 nm (mean ± standard deviation). (**E** to **G**) Testing if *δ* represents a realistic estimate of the localisation accuracy by performing localisation analysis on artificial computer-simulated images (see Methods). (E, F) A representative simulation. Purple point in (F) depicts the real position of the simulated event. (G) Dependency of the localisation precision (defined as the distance to the real position either from the ‘original fit’ or from the ‘averaged position’) on *δ* (N = 1500 simulations, shaded areas depict standard deviation). The simulations reveal that the averaged position determined from the ‘added noise’ stack provides a more reliable estimate for the location of release site. Therefore, we used this improved averaged fit in the analysis of the relative distributions of synchronous and asynchronous release events. Considering that *δ_menn_* = 32.4, the average accuracy in the localisation of exocytosis events in our experimetns was ~ 75 nm (in the range between 50 – 100 nm). (**H**) Comparison of the original and the improved localisation maps for the bouton shown in Fig. 3A-D.

**Fig. S7.**
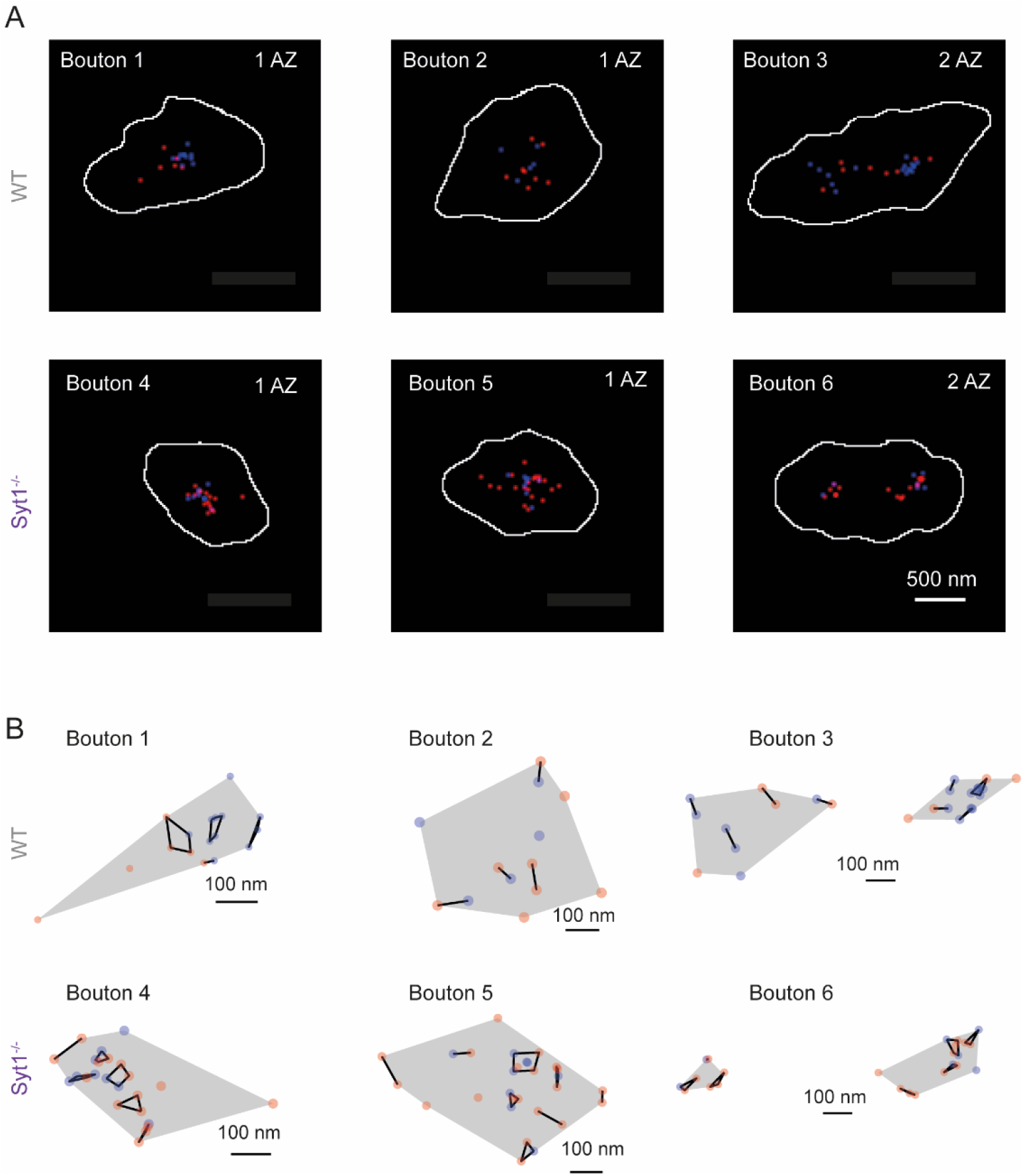
Additional examples of spatial distributions of synchronous and asynchronous exocytosis sites. (**A**) Composite images showing relative locations of synchronous (blue) and asynchronous (red) release events in several representative wild type boutons (WT, 1-3) and Syt1^-/-^ boutons (4 – 6), see also Fig. 3D. (**B**) Hierarchical cluster analysis of vesicular exocytosis sites in the same boutons as in (A). Putative active zones (grey shaded areas) were identified using a clustering diameter threshold of 700 nm. Boutons with 2 active zones (*e.g*. boutons 3 and 6) were excluded from further analysis. To compare the relative locations of synchronous and asynchronous release sites, the positions of vesicular release events were grouped into clusters using a clustering diameter threshold of 100 nm (see Fig. 3F and the main text for details).

## Movie S1. Identification of active presynaptic boutons.

Animated version of Fig. 1B-D.

## Movie S2. Sub-pixel localisation of synchronous and asynchronous release events.

Animated version of Fig. 3C. Scale bar 500 nm.

## References

1. Kaeser, P.S. & Regehr, W.G. Molecular mechanisms for synchronous, asynchronous, and spontaneous neurotransmitter release. Annu. Rev. Physiol. 76:333–63. (2014).

2. Tang, A.H. et al. A trans-synaptic nanocolumn aligns neurotransmitter release to receptors. Nature. 536, 210–214 (2016).

3. Brockmann, M.M. et al. Functional architecture of the synaptic transducers at a central glutamatergic synapse. BioRxiv2020 (2020).

4. Li, S. et al. Asynchronous release sites align with NMDA receptors in mouse hippocampal synapses. Nat. Commun. 12, 677 (2021).

5. Goda, Y. & Stevens, C.F. Two components of transmitter release at a central synapse. Proc. Natl. Acad. Sci. U. S. A. %20;91, 12942–12946 (1994).

6. Huson, V. & Regehr, W.G. Diverse roles of Synaptotagmin-7 in regulating vesicle fusion. Curr. Opin. Neurobiol. 63:42–52. doi: 10.1016/j.conb.2020.02.006., 42-52 (2020).

7. Deng, S. et al. Regulation of Recurrent Inhibition by Asynchronous Glutamate Release in Neocortex. Neuron. 105, 522–533 (2020).

8. Li, J., Deng, S., He, Q., Ke, W., & Shu, Y. Asynchronous Glutamate Release at Autapses Regulates Spike Reliability and Precision in Mouse Neocortical Pyramidal Cells. Cereb. Cortex.bhaa361 (2020).

9. Luo, F. & Sudhof, T.C. Synaptotagmin-7-Mediated Asynchronous Release Boosts High-Fidelity Synchronous Transmission at a Central Synapse. Neuron. 94, 826–839 (2017).

10. Hefft, S. & Jonas, P. Asynchronous GABA release generates long-lasting inhibition at a hippocampal interneuron-principal neuron synapse. Nat. Neurosci. 8, 1319–1328 (2005).

11. Turecek, J. & Regehr, W.G. Neuronal Regulation of Fast Synaptotagmin Isoforms Controls the Relative Contributions of Synchronous and Asynchronous Release. Neuron. 101, 938–949 (2019).

12. Kusick, G.F. et al. Synaptic vesicles transiently dock to refill release sites. Nat. Neurosci. 23, 1329–1338 (2020).

13. Tagliatti, E. et al. Synaptotagmin 1 oligomers clamp and regulate different modes of neurotransmitter release. Proc. Natl. Acad. Sci. U. S. A. 117, 3819–3827 (2020).

14. Marvin, J.S. et al. Stability, affinity, and chromatic variants of the glutamate sensor iGluSnFR. Nat. Methods. 15, 936–939 (2018).

15. Jensen, T.P. et al. Multiplex imaging relates quantal glutamate release to presynaptic Ca(2+) homeostasis at multiple synapses in situ. Nat. Commun. 10, 1414–09216 (2019).

16. Durst, C.D. et al. High-speed imaging of glutamate release with genetically encoded sensors. Nat. Protoc. 14, 1401–1424 (2019).

17. James, B., Darnet, L., Moya-Diaz, J., Seibel, S.H., & Lagnado, L. An amplitude code transmits information at a visual synapse. Nat. Neurosci. 22, 1140–1147 (2019).

18. Geppert, M. et al. Synaptotagmin I: a major Ca2+ sensor for transmitter release at a central synapse. Cell. 79, 717–727 (1994).

19. Sabater, V.G., Rigby, M., & Burrone, J. Voltage-gated potassium channels ensure action potential shape fidelity in distal axons. BioRxiv2020 (2020).

20. Ermolyuk, Y.S. et al. Independent Regulation of Basal Neurotransmitter Release Efficacy by Variable Ca<sup>2+</sup> Influx and Bouton Size at Small Central Synapses. PLoS Biol 10, e1001396 (2012).

21. Maschi, D. & Klyachko, V.A. Spatiotemporal Regulation of Synaptic Vesicle Fusion Sites in Central Synapses. Neuron. 94, 65–73 (2017).

22. Rozov, A., Burnashev, N., Sakmann, B., & Neher, E. Transmitter release modulation by intracellular Ca2+ buffers in facilitating and depressing nerve terminals of pyramidal cells in layer 2/3 of the rat neocortex indicates a target cell-specific difference in presynaptic calcium dynamics. J. Physiol. 531, 807–826 (2001).

23. Grauel, M.K. et al. RIM-binding protein 2 regulates release probability by finetuning calcium channel localization at murine hippocampal synapses. Proc. Natl. Acad. Sci. U. S. A 113, 11615–11620 (2016).

24. Kaeser, P.S. et al. RIM Proteins Tether Ca(2+) Channels to Presynaptic Active Zones via a Direct PDZ-Domain Interaction. Cell 144, 282–295 (2011).

25. Markram, H., Wang, Y., & Tsodyks, M. Differential signaling via the same axon of neocortical pyramidal neurons. Proc. Natl. Acad. Sci. U. S. A. 95, 5323–5328 (1998).

26. Reyes, A. et al. Target-cell-specific facilitation and depression in neocortical circuits. Nat. Neurosci. 1, 279–285 (1998).

27. Edelstein, A.D. et al. Advanced methods of microscope control using muManager software. J. Biol. Methods. 1, 10 (2014).

28. Schneider, C.A., Rasband, W.S., & Eliceiri, K.W. NIH Image to ImageJ: 25 years of image analysis. Nat. Methods. 9, 671–675 (2012).

29. Pernia-Andrade, A.J. et al. A deconvolution-based method with high sensitivity and temporal resolution for detection of spontaneous synaptic currents in vitro and in vivo. Biophys. J. 103, 1429–1439 (2012).

30. Ovesny, M., Krizek, P., Borkovec, J., Svindrych, Z., & Hagen, G.M. ThunderSTORM: a comprehensive ImageJ plug-in for PALM and STORM data analysis and super-resolution imaging. Bioinformatics. 30, 2389–2390 (2014).

31. Rebola, N. et al. Distinct Nanoscale Calcium Channel and Synaptic Vesicle Topographies Contribute to the Diversity of Synaptic Function. Neuron. %20;104, 693–710 (2019).

32. Holderith, N. et al. Release probability of hippocampal glutamatergic terminals scales with the size of the active zone. Nat. Neurosci. 15, 988–997 (2012).

33. Timofeeva, Y. & Volynski, K.E. Calmodulin as a major calcium buffer shaping vesicular release and short-term synaptic plasticity: facilitation through buffer dislocation. Front Cell Neurosci. 9:239. doi: 10.3389/fncel.2015.00239. eCollection@2015., 239 (2015).

34. Ermolyuk, Y.S. et al. Differential triggering of spontaneous glutamate release by P/Q-, N- and R-type Ca(2+) channels. Nat. Neurosci. 16, 1754–1763 (2013).

35. Chamberland, S., Timofeeva, Y., Evstratova, A., Volynski, K., & Toth, K. Action potential counting at giant mossy fiber terminals gates information transfer in the hippocampus. Proc. Natl. Acad. Sci. U. S. A. 115, 7434–7439 (2018).

36. Lou, X., Scheuss, V., & Schneggenburger, R. Allosteric modulation of the presynaptic Ca2+ sensor for vesicle fusion. Nature. 435, 497–501 (2005).

